# Decades-long elevation of interferon-α drives a Sjögren disease endotype: an interdisciplinary study

**DOI:** 10.1101/2025.08.06.668845

**Authors:** Deborah Forbes, Ivana Jorgacevic, Jessica Tarn, Katy R Reid, Bastien Rioux, Kyle Thompson, Kilian Kleemann, Sarah McGlasson, John Casement, Joe Berry, Patrick Müller, Hannah Eberle-Reece, Christa Haase, Brendan Conn, Louis Boon, Karina McDade, William Whiteley, Axel Roers, Wan-Fai Ng, Rayk Behrendt, David Hunt

**Author notes:** Denote equal contributions (Deborah Forbes, Ivana Jorgacevic, Jessica Tarn, Wan-Fai Ng, Rayk Behrendt, David Hunt).

## Abstract

**BACKGROUND:** Mechanistic heterogeneity is a major obstacle to the development of effective treatment for Sjögren disease (SjD) and there is a pressing need to stratify SjD according to precision medicine principles. Aberrant activation of the type I interferon (IFN) pathway represents a leading candidate pathway, but a causal role of elevated IFN-α in driving a Sjögren disease endotype remains to be established.

**METHODS:** We used ultrasensitive single molecule ELISA, and an oligoprotein interferon signature score (derived from broad capture proteomics), to study the role of IFN-α in Sjögren disease. We analysed samples from the UK Primary Sjögren Syndrome Registry (UKPSSR, n=177) and UK Biobank Plasma Proteomics Project (n=47606 without Sjögren, n=257 with Sjögren, including 137 individuals sampled prior to diagnosis) to determine the timecourse and immune endotype associated with elevated IFN-α. To address causality we created a new transgenic mouse model of IFN-α overexpression to establish whether chronically elevated IFN-α drives this immune endotype.

**FINDINGS:** Oligoprotein interferon signatures can be detected at least 14 years prior to diagnosis of Sjögren disease in the UK Biobank-PPP. IFN-α concentrations are elevated in 60% of Sjögren disease patients in the UKPSSR. Individuals with elevated IFN-α display a distinct immunological endotype characterised by cytopenias, hypergammaglobulinaemia, multiple autoantibodies and autoimmunity against the Sjögren autoantigen TRIM21/Ro52. To address the key question of causal direction, we created a new mouse model of systemic chronic IFN-α elevation, in which *Ifn*α*4* is overexpressed by conventional dendritic cells. This model recapitulates key features of the endotype and can be partially reversed by IFNAR1 blockade.

**INTERPRETATION:** Elevation of IFN-α drives an immune endotype of Sjögren disease, originating over a decade prior to diagnosis. SjD patients with elevated IFN-α concentrations are broadly clinically similar to those with normal IFN-α concentrations, yet are immunologically distinct. This highlights the mechanistic heterogeneity of SjD and the need for immunological stratification along precision medicine principles, using high resolution biomarkers. As well as demonstrating causal direction, biological modelling shows that chronic IFN-α elevation over the lifecourse has the potential to establish persistent immune dysregulation which responds only partially to interferon receptor blockade. These findings provide insights into SjD and other “interferonopathic” rheumatological disorders.

**Research in context:** *Evidence before this study:* We searched MEDLINE for “Sjögren’s Syndrome/Disease” and “interferon”, including the terms “subsets”, “sub-groups”, “phenotypes”, and “endotypes”, filtering by “clinical trial”, “stratification”, and “immune-mediated inflammatory”. We also included major review articles from noted experts. We identified reports of associations between IFN-α and SjD, usually using indirect or imprecise measures of IFN-α. None of these studies included prediagnostic samples and causal inference was limited.

*Added value of this study:* This study shows that IFN-α, when measured directly using ultrasensitive single molecule ELISA approaches (uniquely optimised to determine healthy control concentrations), is elevated in a subset of people with SjD with a specific immunological endotype. Analysis of prediagnostic proteomics shows this elevation can be detected up to 14 years before diagnosis. We show that IFN-α drives this endotype (as opposed to vice versa) by recapitulating key endotype features in a novel and unambiguous experimental mouse model of chronic IFN-α elevation. We also show that the pathogenic consequences of IFN-α elevation over long periods of time can only be partially reversed using IFNAR blockade.

*Implications of all the available evidence:* These data have important implications for future research, clinical practice, trial design, and therapeutic development. First, our findings provide clinical evidence, supported by unambiguous preclinical evidence, that decades-long elevated IFNα can cause and drive a SjD endotype – and accurately defines the level of heterogeneity. Secondly, we provide biomarkers which may be of use in stratifying clinical trial design and also for early identification of at-risk individuals. Thirdly we provide biological proof of principle that longstanding and potentially undiagnosed elevation of IFN-α can establish persistent immune dysregulation which may respond only partially to IFNAR blockade. Together these findings inform precision medicine approaches and future trial design.

## INTRODUCTION

Sjögren disease (SjD) is a systemic autoimmune condition which affects exocrine glands and is associated with specific immunological abnormalities such as autoantibodies against Ro/La antigens ^1^. Despite several clinical trials of systemic immunosuppression in SjD, no disease modifying treatments have been proven to be effective ^2^. An insufficient understanding of immune mechanisms which drive disease – and their heterogeneity - is a contributory factor to failed trials ^3^. As such it is important to identify dysregulated immunological pathways which drive disease and identify subpopulations who may respond to targeting these pathways ^4^. The identification of such immunopathological endotypes is a priority of precision medicine ^5^ ^6^ ^7^ ^8^.

Within the framework of precision medicine, “stratified medicine” is defined as “The use of a molecular assay to define subpopulations, rather than individuals, who are likely to benefit from a treatment intervention” ^9^. The type I interferon pathway is an important candidate pathway to evaluate as a stratification tool in SjD. Type I interferons, which include 13 IFN-α subtypes, are pleiotropic cytokines and act via the common type I interferon receptor IFNAR, leading to upregulation of hundreds of interferon stimulated genes ^10^. Activation of the type I interferon response has been identified in a number of autoimmune diseases, most notably systemic lupus erythematosus (SLE) where 80-90% of patients have evidence of a transcriptomic interferon signature ^11^ ^12^, although the degree to which type I interferon proteins drive disease is less clear. This is highlighted by the variability of clinical response to interferon receptor blocking drugs observed in phase 3 clinical trials in SLE ^13^ ^14^. Cohort studies have identified elevated IFN-α and interferon stimulated gene expression signatures in some SjD patients ^15^ ^3^ ^16^.

A key unresolved question is whether elevated IFN-α drives aspects of immune disease in SjD or is simply an epiphenomenon, itself driven by underlying immune dysregulation in certain patients. To date there have been three limitations on resolving this question. Firstly, precision medicine requires precision tools, which have been lacking for the quantification of IFN-α concentrations in human samples. IFN-α proteins are low abundance and highly potent proteins, and the tools for accurate measurement of IFN-α concentrations in human patient samples have been largely limited to indirect measures, because peripheral type I IFN protein concentrations are below the limit of detection of conventional ELISA technologies ^17^ ^12^. Secondly, little is known about the chronicity of IFN-α elevation and how this relates to disease onset. This is because most traditional disease cohorts do not have pre-diagnosis samples. Resolving this question is important not only for understanding if IFN-α drives disease, but could also provide practical approaches for identification of at-risk individuals. Thirdly, there are few, if any, viable mouse models of chronic IFN-α elevation which allow unequivocal causal inference to be drawn about the downstream immunological consequences of chronic type I IFN activation.

Therefore to address this key question we used emerging protein detection technologies (ultrasensitive single molecule ELISA and Olink proteomics) to define an IFN-α associated endotype of SjD in 60% of patients, and show this cytokine elevation arises over a decade prior to diagnosis in UK Biobank. Core clinical features of this endotype include blood cytopenias, hypergammaglobulinaemia and autoimmunity directed against the Sjögren-associated autoantigen TRIM21. To demonstrate that these clinical features are driven by chronic IFN-α elevation, we recapitulate this endotype in a novel mouse model of chronic interferon-driven disease, achieved through overexpression of *Ifn*α*4* in conventional dendritic cells type 1. We also show that some, but not all, aspects of this endotype can be reversed with interferon receptor blockade in this model. Together, these findings provide a framework and practical tools for precision medicine approaches in SjD.

## METHODS

### The UK Primary Sjögren’s Syndrome Registry (UKPSSR)

The UKPSSR is a national observational cohort of clinically well characterised patients with SjD, who fulfil the 2002 American European Consensus Group classification criteria ^18^ and were recruited from 40 UK based centres. All patients recruited to the UKPSSR had peripheral blood, serum, DNA, and RNA banked at the time of recruitment. All samples were stored in Eppendorf tubes at −80°C until they were used in our analyses ^19^. The ESSDAI questionnaire was scored by a member of trained clinical staff on the same date.

The registry recruited healthy controls at the same time as SjD patients. These are non-SjD individuals, with no history of fatigue, dry eyes/mouth or autoimmune disease who were age matched to ±3 years of the patient group.

All participants gave written consent for collection of clinical data and peripheral blood samples at the time of recruitment to the cohort. Research ethics approval was granted by the UK National Research Ethics Committee NorthWest – Haydock.

A group of 177 SjD patients and 28 healthy controls (plus a further eight healthy controls from the Future MS study, see supplementary methods) from the UKPSSR were chosen for this study. Their selection was based on the availability of previous microarray and Olink proteomic data ^20,21^.

### UK Biobank and Pharma Proteomics Project

The UK Biobank (UKB) is a large population-based cohort of 502,000 participants aged 40-69 years old during the period of recruitment from 2006 and 2010. At baseline, details of current medical conditions were collected through a self-reported questionnaire and a structured interview alongside blood, urine, and saliva for future biological phenotyping. During follow-up health related outcomes are collected through data linkage with hospital records, national death registries, and primary care records. Diagnoses are recorded as both Read (v2 and v3) codes from primary care records and ICD (v9 and v10) codes from hospital admission and death records ^22^.

The UKB Pharma Proteomics Project (UKB-PPP) used Olink Explore 3072 to measure the plasma concentration of 2923 unique proteins in 54219 UKB participants. This subset of UKB participants was comprised of 46595 randomly selected participants and 6376 participants selected by UKB-PPP consortium members.

The UK Biobank (UKB) has received ethical approval from the North West Multicentre Research Ethics Committee, and all participants provided written informed consent at recruitment. This research was conducted under the UKB Resource application number 93160.

### Definition of cases in the UK Biobank

A case of SjD was defined as at least one diagnostic code from inpatient hospital, primary care or death records, or any self-reported diagnosis at any assessment visit, with codes detailed in supplementary methods. The definition of SjD cases using administrative health data is commonly used and has a high specificity (>90%) in healthcare systems with universal coverage using ICD codes from hospitalizations or physician claims. ^23^.

### Whole blood transcriptomics

Whole blood was collected in PAXgene tubes from UKPSSR participants and transcriptomics performed as previously described. A stepwise (forward) multiple linear regression filter was used to identify which of the 144 interferon stimulated gene transcripts from Chaussabel modules 1.2, 3.4, and 5.12 were most strongly associated with blood IFN-α concentration. Data were split into training and testing sets (training n = 137, testing n = 67) and the maximum R^2^ for the validation set was used as the stopping rule for variable selection. This analysis was performed in JMP Pro V.15 a.

### Quantification of serum IFN-**α** protein using a pan-IFN-**α** Simoa

Serum IFN-α was measured in the UKPSSR with a prototype pan IFN-α single molecule array digital ELISA (Simoa) following the manufacturer’s instructions (Quanterix, USA). This was performed on-the HD-X Analyser platform (Quanterix, USA, serial no. 2710020203, Software version 3.0.2003.4001). Samples were thawed on ice, then loaded neat and subject to a 2-step assay with a one in two on-board dilution with sample diluent to give two replicates per sample. There were no helper beads. The HD-X analyser calculated a calibration curve based upon triplicate measurements of recombinant IFN-α17 and used this to determine the concentration of IFN-α in each sample. All measurements were performed blinded to clinical details.

### Olink proteomics

Olink is a high throughput protein biomarker discovery platform that employs a proximity extension assay to measure the relative concentrations of up to 2932 unique analytes per sample. In brief, there are two paired antibodies per analyte which bind simultaneously to their target protein. Each antibody pair has a unique DNA oligonucleotide label, when the antibodies are bound and in close proximity, these oligonucleotides hybridize and are extended by DNA polymerase. This generates a unique DNA code for each protein, which is amplified by PCR, read out by NGS or qPCR, and normalised using a log2 transformation to give a normalised protein expression (NPX) for each analyte.

### Olink proteomics in the UK Primary Sjögren syndrome Registry

Anonymised samples for 39 SjD patients were sent to Olink proteomics for quantification of 454 proteins across five multi-analyte Target 96 panels, ‘Inflammation’, ‘Immune Response’, ‘Organ Damage’, ‘Cardiovascular III’ and ‘Metabolism’. This subset of patients were selected randomly and to have a balance of symptom based phenotypes ^3,21^. To remove likely nuisance proteins and prevent overfitting and unstable coefficients we first rationalized the number of proteins included in our model by referring Chaussabel modules M1.2, M3.4 and M5.12 to identify and include known ISGs only represented in Olink (five Target 96 panels in UKPSSR and Explore 3072 in UK Biobank; n=9). Then we employed statistical variable selection from univariable linear regressions to include five proteins with a positive association with log-IFN-α at p<0.1. We did not exclude non-secreted proteins as they may be detectable in the serum through leakage from cells that are either active or undergoing cell death ^24^. From these five proteins, an oligoprotein score, which we termed Sjögren Interferon-α Response in Olink (SIRO) score, was generated as a proxy for IFN-α measurement that could be applied to both the UKPSSR and to the UKB.

### Olink proteomics in the UK Biobank

The UKB Pharma Proteomics Project used Olink Explore 3072 to measure the plasma concentration of 2923 unique proteins in 54219 UKB participants. This subset of UKB participants was comprised of 46595 randomly selected participants, 6376 participants selected by UKB-PPP consortium members, and 1268 participants who participated in the COVID-19 repeat imaging study. For our analysis we excluded the 1268 participants who were part of the COVID repeat imaging study. Participants were included in our analysis if they had complete data for all five SIRO proteins (n=47,863; mean=0, SD=0.60). There were 257 UKB participants with a diagnosis of SjD who were part of the UKB-PPP with complete data.

### Clinical laboratory results

Blood samples were taken from all participants on the day of recruitment in EDTA tubes. On the same day these were transferred to the pathology lab of the recruiting NHS hospital where the standard local procedure for determining full blood counts and immunoglobulin concentrations were used.

### Autoantibody testing

A subset of SjD patients (n=140) had autoantibody testing performed at the NHS Gateshead Regional Laboratory at the time of entry into the UKPSSR cohort. These results were used to classify patients into seronegative or seropositive for the clinically relevant antibodies to antinuclear (ANA), mitochondrial (AMA), thyroid peroxidase (ATPO), citrullinated c-peptide (CCP), centromere (CENT), gastric parietal cells (GPC), double stranded DNA (ds-DNA), Jo1 smooth muscle (SMA), rheumatoid factor (RHF), ribonuclear protein (RNP), SSA/Ro (RO), SSB/La (LA), Scl70 (SCL70), and smith (SM).

In addition, 127 patients’ sera were tested for fine specificity SSA antibodies to the TRIM21/Ro52 epitope, with 7ug/mL as the upper limit of normal. These tests were performed following the standard protocol for the NHS Service.

### Cox Regression

We used Cox proportional hazards models to test the association of TRIM21 strata with SjD, multiple sclerosis or haemorrhoids in three separate models. We used two controls. Firstly, we used haemorrhoids as negative control outcome because it is a common condition which is unlikely to be related to type I interferons. Further this condition is subject to similar patterns of case ascertainment using administrative health databases. We have also used multiple sclerosis as a negative control since this is an autoimmune disease where our single molecule ELISA studies have previously shown a lack of association with IFN-α ^25^. Analyses were adjusted for potential predictors of autoimmunity, including age, sex, self-reported ethnic background, smoking (current, previous or never), Townsend deprivation index and presence of other autoimmune disease. Participants were excluded if they reported being on interferon therapy (n=9) or if they had missing data on any of the covariates (n=315). Participants were followed from baseline until the first of outcome occurrence, death, loss of follow-up, or study end date (October 31, 2022). The study end date was defined as the last day of the month with mostly complete data (as per the UKB definition) for the main source of diagnoses (National Health Service [NHS] England inpatient data).

### Statistical analysis

Statistical analysis and graphing of the single-molecule ELISA and clinical data were performed on graphpad prism software and R statistical programming software. Spearman’s Test was used to assess correlation between tests. To assess significance between two variables, the non-parametric Mann-Whitney U test was employed. Analysis of the transcriptomic arrays was performed using Bioconductor libraries in the R environment for statistical computing ^26^. Statistical analyses in UKB were conducted on the Research Analysis Platform using JupyterLab (v3.6.1) and R (v4.2.0) with packages UpSetR (v1.4.0), ggplot2 (v.3.4.4) and survival (v3.7.0).

### Methods for transgenic mice

Mouse interferon alpha 4(Ifna4) cDNA (ENSMUST00000094973.5) was cloned via AscI restriction site into the Rosa26-specific pSERC plasmid (kind gift from Thomas Wunderlich, MPI for Metabolism Research, Cologne) resulting in the following configuration from 5’-3’: 5’arm of homology, CAGs promoter, Lox-STOP-Lox cassette (NEO resistance), Ifna4 cDNA, IRES EGFP, 5’arm of homology, PGK-driven diphtheria toxin negative selection marker. Mice were generated by electroporation of the linearized targeting construct into Agouti JM8A1.N3 mouse embryonic stem cells resulting in 14 neomycin-resistant ES cell clones, of which 10 had integrated the full-length construct as assesses by long range PCR and southern blot for both homology arms. One clone was injected into C57BL6/N morulae and germline transmission was achieved to establish the Rosa26-mIFN`U`4 (“R26-mIFN`U`4”) mouse line (Figure 5A). To achieve deletion of the STOP cassette only in conventional dendritic cells, R26-mIFN`U`4 mice were crossed to Clec9a-Cre mice (Kind gift from Caetano Reis e Sousa, The Francis Crick Institute, London) ^27^. For the treatment approach, anti-IFNAR1 antibody (clone: MAR1-5A3, Biolegend) or Isotype (clone: MOPC-21, Biolegend) in the concentration of 20mg/kg was injected 3 times per week i.p, starting at 4 weeks age or 250 μg rat anti-mouse– CD8β antibody (Ab) (clone YTS156.7.7, referred as anti-CD8β Ab) or isotype-matched irrelevant rat-anti-Phyt1 (clone AFRC-MAC51, referred to as a Isotype).

All animals were bred at Experimental Center of the Medical Faculty at the University Hospital Dresden and at the Haus für Experimentelle Therapie (HET), University Hospital Bonn under the approved licenses TVV 54/2017, 88/2017 (both Landesdirektion Sachsen) and 81-02.04.2020.A037, 81-02.04.2023.A259 (both LANUV, NRW) respectively. For targeted analysis female and male mice were used interchangeably and sacrificed at the age of 6 weeks.

Further detailed mouse methods, including statistical approaches, are outlined in supplementary methods.

## RESULTS

### IFN-***α*** is elevated in a subpopulation of Sjögren disease and drives the transcriptomic response

We measured IFN-α serum concentrations in a well-characterised cohort of patients with SjD using an optimised single molecule ELISA assay (Figure S1A-C). This assay uses a highly specific, high-affinity pair of antibodies originally isolated from patients with autoimmune polyendocrine syndrome type I to reproducibly quantify multiple subtypes of IFN-α in the attomolar range ^12^. This unprecedented sensivity enables the quantification of IFN-α concentrations even within healthy controls and non-interferonopathic disease ^12^. IFN-α concentrations were significantly higher in the UK Primary Sjögren Syndrome Registry cohort, (UKPSSR, Figure 1A) (n = 177) compared to healthy controls (n = 36) (Figure 1B) and 61% of SjD patients with serum concentrations of IFN-α above the healthy control range (Figure 1B). Paired IFN-α Simoa and transcriptomic data showed strong correlation between IFN-α concentration and transcriptomic evidence of type I IFN activation (Figure 1C, S1D-E), suggesting this signature is primarily driven by IFN-α in people with SjD.

**Figure 1.**
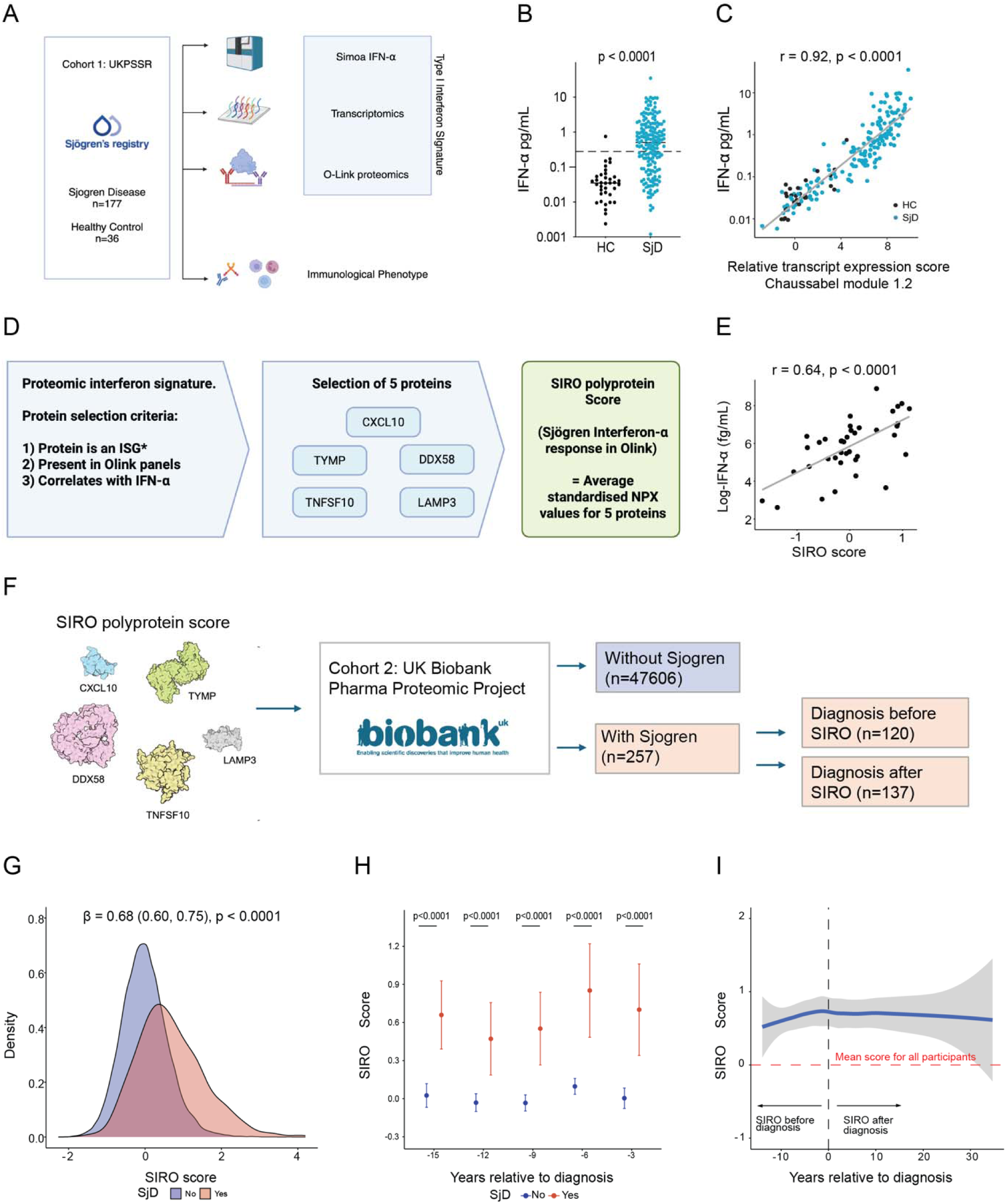
Ultrasensitive IFN-α detection and development of an interferon polyprotein score reveals interferon signatures in patients with SjD over a decade prior to diagnosis. (A) Overview of the SjD cohort for paired Simoa studies in the UK Primary Sjögren’s Syndrome Registry (UKPSSR). Serum samples were taken from SjD patients enrolled in the UKPSSR (n=177) and healthy controls (n=36) and serum IFN-α was quantified using a pan-IFN-α Simoa, and compared to transcriptomic and proteomic analyses. (B) Patients with SjD have higher IFN-α than healthy controls (median=0.5032 (IQR 0.1053– 1.549) pg/mL vs 0.03481 (IQR 0.01651–0.05359) pg/mL, Mann Whitney U test p<0.0001). The upper limit of normal serum IFN-α was defined as mean log(IFN-α) plus 1.96SD in healthy controls (0.279pg/mL, represented by dashed line). SjD patients were then classified as having normal IFN-α (39%) or elevated IFN-α (61%). (C) Serum IFN-α is associated with the type I IFN transcriptomic response. A microarray for the 23 ISGs in Chaussabel modules M1.2 was used to quantify the transcriptomic response to IFN in the serum of 177 SjD patients and 28 HCs. Each individual was given a score relative to the mean HC expression of all trancripts in module M1.2. This correlated with Simoa IFN-α (Spearman’s rank r=0.89, p<0.0001). (D) Overview of the approach to develop the polyprotein interferon score. Olink proteomic analysis was performed on a subset of UKPSSR SjD patients (n=39), univariable linear regressions were applied to select proteins with a positive association with IFN-α at p-value <0.1, and these were used to create the Sjögren Interferon-α Response in Olink for Sjögren disease (SIRO) score, a correlate of IFN-α activity that could be applied to both UKPSSR and UKB to explore the relationship between IFN-α activity and the diagnosis of SjD. *ISGs defined by Chaussabel modules 1.2, 3.4, 5.12. (E) Serum IFN-α is associated with a polyprotein marker of IFN-α response (SIRO score) in UKPSSR participants (β=1.39; 95% CI: 0.83, 1.95; p<0.0001; adjusted R-squared=38.8%; Pearson’s correlation coefficient=0.64, p<0.0001). (F) Summary of approach to detecting polyprotein interferon signatures in UK Biobank using SIRO. The UKB Pharma Proteomics Project used the Olink 3072 panel to quantify serum protein concentrations for over ∼54,000 participants’ baseline samples. We included all participants who had complete data for the five SIRO proteins (without SjD: n =47606; with SjD: n=257). (G) The SIRO score is significantly higher in UK Biobank participants with SjD than those without (mean=0.67, SD=0.86 vs. mean=-0.01, SD=0.60; β=0.68; 95% CI: 0.60, 0.75; p<0.0001). (H) The SIRO score is elevated over a decade before SjD diagnosis in the UK Biobank. We compared SIRO scores in participants with and without SjD by 3-year strata before diagnosis using a nested case-control design, whereby each participant with SjD was matched with 10 participants without SjD on age at baseline and sex (exact match). The plot presents the mean SIRO score by group along with their 95% CIs and p-values from linear regressions by strata of years before diagnosis. (I) SIRO scores relative to time of SjD diagnosis in UK Biobank. As well as elevation prior to diagnosis, polyprotein interferon scores remained elevated for up to 30 years after diagnosis. “SIRO before diagnosis” refers specifically to datapoints from individuals who had Olink analyses performed before a diagnosis of SjD was recorded in UK Biobank. “SIRO after diagnosis” refers to individuals with Olink analyses performed after diagnosis of SjD. Individual datapoints displayed in Supplementary Figure 2B. Smooth curves with standard errors using the locally estimated scatter plot smoothing (LOESS) method.

### A polyprotein interferon signature in Sjögren disease

We then developed and validated a type I interferon polyprotein signature (Figure 1D). We performed serum proteomic analyses of 39 UKPSSR participants, quantifying 454 proteins across five multi-analyte Target 96 panels (Figure 1D). From a pool of candidate proteins shown to be upregulated ISGs in our transcriptomic analyses, we identified five proteins (CXCL10, TYMP, DDX58, LAMP3, TNFSF10) with a positive association (β >0) and p-value <0.1 in univariable linear regressions with log-IFN-α (fg/mL) (Figure 1D). We then derived a polyprotein score which we termed the “SIRO score” (Sjögren Interferon-α Response in Olink). The SIRO polyprotein score was strongly associated with log-IFN-α (fg/mL) in SjD patients (Figure 1E). To further validate that the SIRO score is strongly related to blood IFN-α concentrations we measured SIRO in UKB participants who were receiving recombinant type I interferon therapy. We found strong elevation of SIRO in these individuals that could not be explained by their underlying diseases (Figure S2C). We also showed that SIRO score in SjD was influenced by HLA-DQA1*05:01 allele dosage, a genetic association known to be associated with increased IFN-α in SjD (Figure S3) ^16^.

### Polyprotein interferon signatures are elevated in patients with Sjögren disease over a decade prior to diagnosis

The UK Biobank Pharma Proteomics Project (UKB-PPP) ^28^ facilitates the study of the plasma proteome of ∼50000 UK Biobank participants, including samples taken many years before a diagnosis of SjD is made (supplementary table 4). We applied our SIRO polyprotein score to UKB-PPP participants with SjD (n=257) and without SjD (n=47606) (Figure 1F, Figure S2A). These participants with SjD had significantly higher SIRO polyprotein scores (mean=0.67, SD=0.86) as compared to those without SjD (Figure 1G). Just over half of participants with SjD had a diagnosis recorded after baseline sampling used for proteomic analysis (n=137; 53.3%). We compared the SIRO polyprotein scores in participants with SjD, versus without SjD by 3-year strata before diagnosis using a nested case-control design (10 controls per case matched on age at baseline blood sampling and sex). We found that SIRO polyprotein scores were consistently increased in people with SjD over a 15-year period prior to diagnosis and 30 years after diagnosis (Figure 1H and 1I, Figure S2B).

### A distinct immune endotype of Sjögren disease is associated with elevated IFN-***α***

We next analysed the relationship between blood IFN-α concentrations and specific, clinically relevant, immunological features of patients with SjD in UKPSSR, with the aim of using IFN-α Simoa to stratify SjD (Figure 2A, Table S2). We found strong associations between IFN-α and peripheral blood cytopenias, notably white cell count (WCC) (Figure 2B), lymphopenia (Figure 2C) and neutropenia (Figure 2D), but not thrombocytopenia (Figure 2E). We identified a strong association between IFN-α and IgG concentrations, as well as IgA concentration (Figure 2F 2G), but no association with IgM concentrations (Figure 2H). We observed an association between IFN-α concentrations and the development of antinuclear antibodies (ANA) and multiple autoantibodies (Figures 2I, 2J and 2K). We found no evidence of association between IFN-α concentrations and disease activity measured by EULAR Sjögren’s Syndrome Disease Activity Index (ESSDAI), pattern of organ involvement, nor anxiety or fatigue (Figure 2L, S1F-H Table S3) (Seror et al., 2015). Therefore Simoa facilitates stratification of patients into subpopulations of SjD with elevated (∼60%) or normal (∼40%) IFN-α, with distinct immunological endotypes (Figure 2A) yet broadly similar patterns of disease activity and organ involvement.

**Figure 2.**
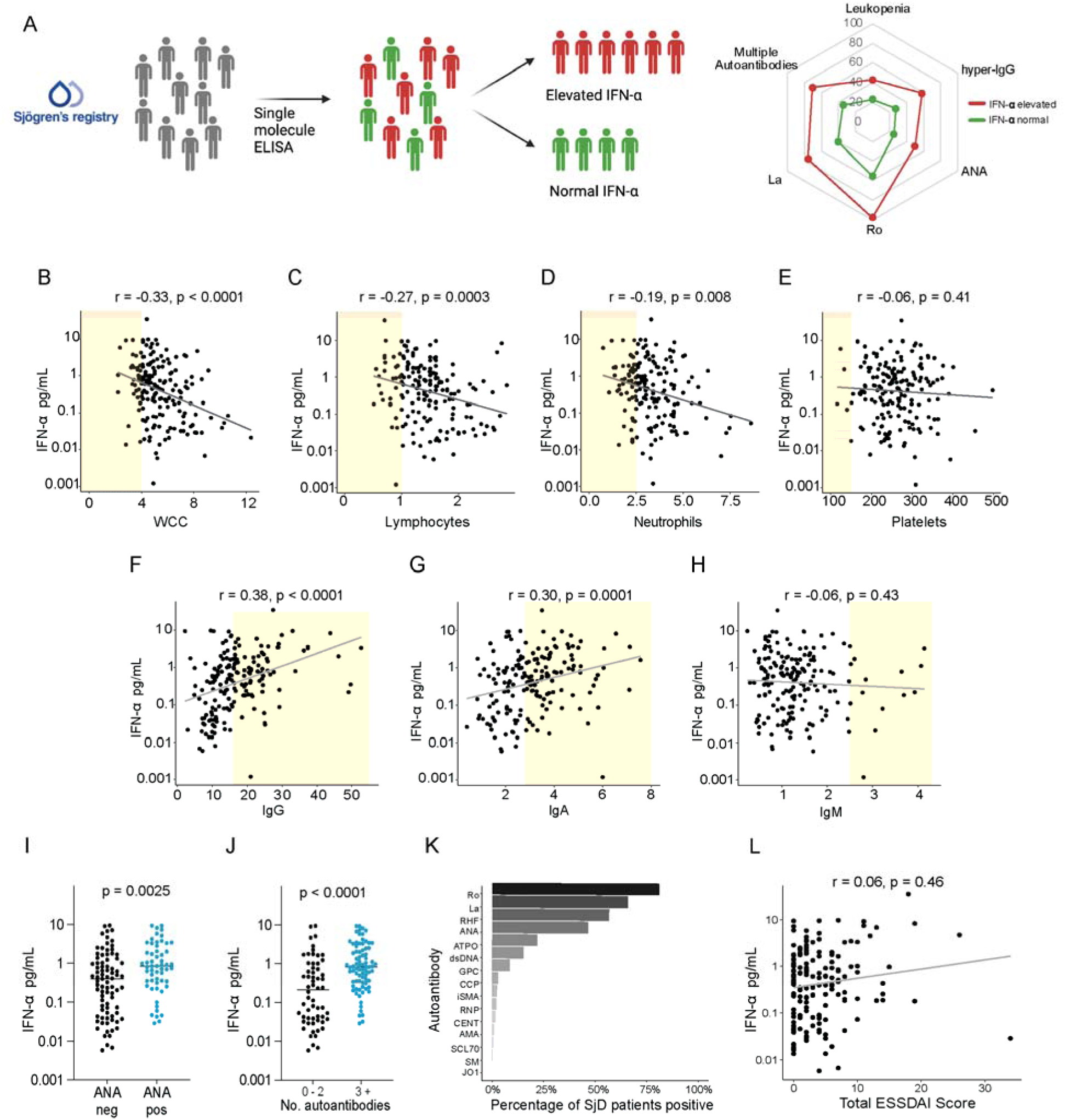
Elevated IFN-α in individuals with Sjögren disease is associated with a distinct immune endotype characterised by white blood cell cytopenias, hypergammaglobulinaemia and autoantibody production. (A) Stratification of the UKPSSR cohort using Simoa IFN-α. The log mean+1.96SD IFN-α of our healthy controls (n=36) was used as the upper limit of normal (0.279pg/mL). Using this cut off we identified SjD patients as having normal (39%, n=69) or elevated (61%, n=108) IFN-α. The proportions (%) of each group with these features is represented as a spider chart here. Using the upper limit of normal for serum IFN-α based upon our controls, SjD patients were stratified into elevated IFN-α and normal IFN-α groups. There was a significant difference between the proportion of these groups with leukopenia, hypergammaglobulinaemia, and antibodies to ANA, Ro, La, and multiple autoantibodies (Defined as greater than three autoantibodies present on the systemic panel at baseline). (B - E) IFN-α and peripheral blood counts. The regional NHS clinical laboratory performed peripheral blood cell counts in 177 SjD patients. IFN-α was negatively correlated with total white blood cell count (B) (r =-0.33, p<0.0001), lymphocyte count (C) (r =0.27, p=0.0003) and neutrophil count (D) (r=-0.19, p=0.008). There was no significant association between IFN-α and platelet count (E) (r=-0.06, p=0.41). The yellow shaded areas represent clinically abnormal results as per the regional NHS laboratory reference range. (F - H) Serological analysis showed that Simoa IFN-α concentration was positively associated with elevated IgG (F) (r=0.38, p<0.0001) and IgA (G) (r=0.30, p<0.0001). There was no significant association between IFN-α and IgM (H) (r=-0.06, p=0.43). (I) SjD patients who are ANA seropositive have higher serum IFN-α than ANA seronegative SjD patients (median IFN-α 0.8431(IQR 0.3227 – 2.436) pg/mL vs. 0.3912 (IQR 0.05343 – 1.229) pg/mL, Mann Whitney U test p=0.0025). (J & K) IFN-α is associated with breadth of autoantibody production in SjD. Patient sera (n =140) was subjected to a standard screening panel for common clinically relevant autoantibodies. (J) Patients with more than three autoantibodies had higher IFN-α than those with 0-2 autoantibodies (Median IFN-α 0.2073 (IQR 0.325 – 2.414) pg/mL vs 0.8150 (IQR 0.03747 – 0.9457) pg/mL Mann-Whitney U, p < 0.0001). (K) There was a broad autoantibody repertoire in the SjD patient group. The rates of seropositivity were as follows 80.9% Ro/SSA, 65.8% La/SSB, 56.6% RhF, 46.5% ANA, 21.9% ATPO, 15.2% dsDNA, 8.42 GPC, 2.89% CCP, 2.41% iSMA, 2.17% RNP, 1.69% Cent, 1.20% AMA, 0.723% SCL70, 0.48% SM, 0% Jo1. (L) Serum IFN-α and clinician reported disease severity have no significant association in the UKPSSR cohort. At the time of recruitment to the cohort and serum sampling, the ESSDAI was reported by the local clinical team. We show that there is no association between serum IFN-α concentration and ESSDAI score (Pearson’s correlation coefficient=0.06, p=0.46). ANA – Anti-nuclear, AMA Anti-mitochondrial, ATPO Anti-thyroidperoxidase, CCP Citrullinated c-peptide, CENT - Centromere, GPC - Gastric parietal cell, dsDNA - Double stranded DNA, iSMA smooth muscle, RhF - rheumatoid factor, RNP - ribonuclear protein, SM – Smith. ESSDAI – EULAR Sjögren Syndrome Disease Activity Index, UKPSSR – UK Primary Sjögren Syndrome Registry.

### IFN-***α*** elevation is associated with autoimmunity against the Sjögren Disease-associated autoantigen TRIM21

TRIM21 is a cytosolic E3 ubiquitin protein ligase and antibody receptor, which is recognised by Sjögren disease-associated Ro52 autoantibodies and thus a key Sjögren autoantigen ^29^ ^30^. Notably, our paired Simoa-transcriptomic analysis in UKPSSR SjD patients (n=177) identified *TRIM21* as one of the most highly upregulated ISGs by IFN-α in SjD (Figure 3A). Strong correlation was observed between IFN-α concentration and *TRIM21* transcript level in peripheral blood (Figure 3B) but there was no correlation with transcript level for other SjD-associated autoantigens *La* and *Ro60/TROVE2* (Figure S1I). IFN-α concentration was significantly higher in TRIM21/Ro52 autoantibody positive individuals, compared to antibody-negative (Figure 3C). We observed a strong negative association between IFN-α and serum TRIM21 concentration quantified by Olink (Figure 3D), and found that low serum TRIM21 concentrations were related to the presence of TRIM21/Ro52 autoantibodies (Figure 3E). We identified UKB-PPP participants with baseline measurement of TRIM21 levels through Olink Explore 3072 (n=51,372)(Figure 3F) and tested the association of low TRIM21 levels with incident diagnoses of SjD and two negative control outcomes (multiple sclerosis, an autoimmune disease where we have shown there is no elevation of IFN-α ^25^ and haemorrhoids) in Cox proportional hazards models. Low TRIM21 levels (<-2SD) at baseline were strongly associated with incident SjD diagnosis (n=145) as compared to the reference (−2SD to +1SD), but not with multiple sclerosis or haemorrhoids (Figure 3G-I). Low serum TRIM21 concentrations are also observed in 17% of people with a diagnosis of SLE in UK Biobank (Figure 3F). We tested whether, in this setting, low TRIM21 levels provided additional information to predict the risk of SjD in people with an established diagnosis of SLE (previously referred to as “secondary” Sjögren Syndrome). In 337 participants with SLE but without SjD at baseline, low TRIM21 levels (< −2SD) were significantly associated with incident SjD diagnosis after adjustment for age, sex, ethnicity, smoking and Townsend deprivation index as compared to the reference (≥-2 SD to ≤1 SD) using a Cox regression. Therefore individuals with SLE and low Olink TRIM21 concentrations were more likely to develop “secondary” SjD in the following decade than those who have normal TRIM21 concentrations (Figure 3J).

**Figure 3.**
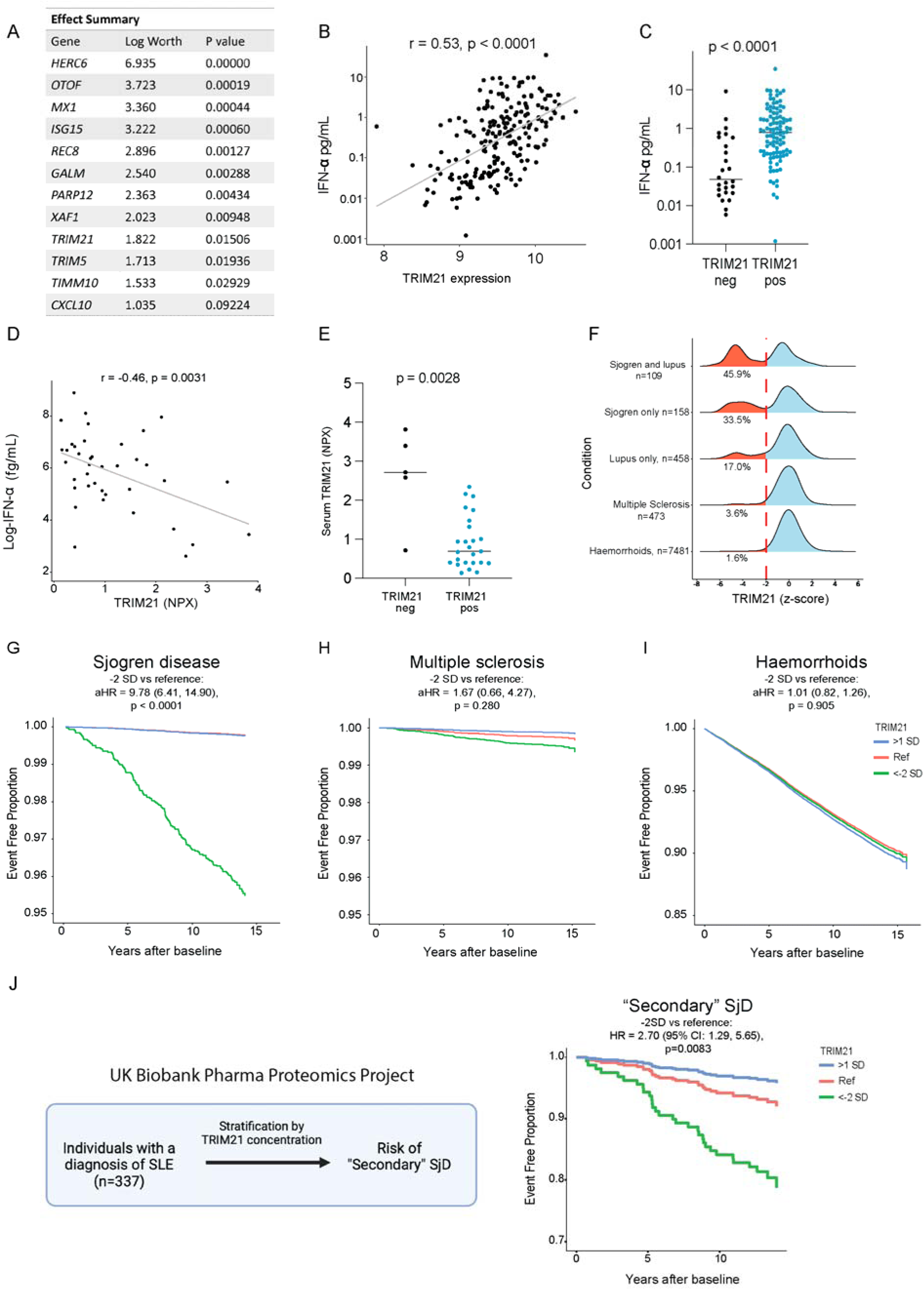
IFN-α elevation is associated with upregulation and autoimmunity against the Sjögren Disease autoantigen TRIM21. (A) Table displaying the top 12 genes whose expression is most strongly associated with blood IFN-α in SjD patients, identified using a stepwise regression filter. *TRIM21* is the ninth most strongly associated gene with IFN-α in our model, and encodes the TRIM21/Ro52 protein which is a target of autoantibodies in SjD. (B) Transcriptomic expression of *TRIM21* was measured using a whole blood microarray in 177 SjD patients. There was a strong positive correlation with serum IFN-α (Spearman’s rank r=0.53, p<0.0001). (C) Fine specificity serostatus for antibodies to the TRIM21/Ro52 epitope was available for 127/177 SjD patients. The cut off for TRIM21/Ro52 antibody positivity was 7 ug/mL. TRIM21/Ro52 seropositive patients (n=101) had higher serum IFN-α than TRIM21/Ro52 seronegative patients (n=26) (Median 0.7824 (IQR 0.233 – 2.028) vs. Median 0.040807 (0.02171 – 0.5875) pg/mL, Mann Whitney U, p<0.0001). (D) Serum IFN-α is strongly negatively associated with serum TRIM21 concentration quantified by Olink (n=39 SjD patients, r = −0.46, p=0.0031). (E) TRIM21/Ro52 seropositivity is associated with lower serum TRIM21 protein levels. Thirty SjD patients had both TRIM21/Ro52 serostatus and Olink proteomic performed. UKPSSR patients with TRIM21/Ro52 antibodies(n=25) had lower TRIM21 protein on Olink analysis than those without TRIM21/Ro52 antibodies (n=5) (Median 2.72 (IQR 1.651 – 3.609) vs. 0.692 (IQR 0.4011 – 2.402) NPX). (F) Serum TRIM21 is reduced in a much higher proportion of UK Biobank participants with SjD (38.7%) than in those with multiple sclerosis (3.6%) or haemorrhoids (1.6%). Low serum TRIM21 concentrations are also identified in 17% of people with SLE, another condition where Ro antibodies can be detected, albeit at lower frequency than SjD. (G)-(I): Low TRIM21 levels at baseline were strongly associated with incident diagnosis of SjD but not MS or haemorrhoids in UKB. We performed a survival analysis to assess the association between standardised TRIM21 levels and incident SjD, multiple sclerosis and haemorrhoids diagnosis. Low TRIM21 was defined as <-2SD from the mean and a Cox proportional hazards model was used to test the association between TRIM21 and diagnosis. Low TRIM21 levels at baseline were strongly associated with incident SjD diagnosis (n=145) with an adjusted hazard ratio (aHR) of 9.78 (95% CI: 6.41, 14.90; p-value <0.0001) as compared to the reference. In contrast, low TRIM21 concentrations (<-2SD) at baseline was not associated with incident multiple sclerosis (n=120, aHR=1.67, 95% CI: 0.66, 4.27; p=0.280) nor with haemorrhoid diagnosis (n=4,197, aHR=1.01, 95% CI: 0.82, 1.26; p=0.905). Note: ‘survival rate’ refers specifically to event-free survival rate. (J) Participants in UK Biobank with a diagnosis of SLE with reduced serum TRIM21 levels were significantly more likely to develop a diagnosis of SjD (previously referred to as “secondary Sjögren Syndrome”) than those who had normal TRIM21 concentrations. Adjusted curves for incident SjD in people with SLE by TRIM21 levels at baseline in UK Biobank (low vs reference) using a Cox regression adjusted for age, sex, ethnicity, smoking and Townsend deprivation index with HR=2.70 (95% CI: 1.29, 5.65; p=0.0083).

Therefore, our analyses of patients with SjD show that chronic activation of the type I interferon response, which occurs over a decade before clinical manifestations and diagnosis, is associated with a specific immune endotype (Figure 2A) characterised by peripheral blood cytopenias, immunoglobulin and autoantibody production, including autoreactivity against TRIM21, an important autoantigen in SjD.

A fundamental question persists as to whether chronic elevation of IFN-α drives this immune endotype. Causation is difficult to demonstrate in observational human studies since IFN-α elevation may itself simply reflect a response to wider immune dysregulation. We therefore looked to an animal model system to help overcome these limitations on causal inference.

### A transgenic model of chronic IFN-***α*** overexpression

We considered which models might provide unambiguous insights into the relationship between IFN-α and the associated immunological phenotypes we observed in SjD. A number of animal models for SjD do exist, some of which exhibit elevation of IFN-α ^31^ ^32^. However, teasing out the contribution of IFN-α in the context of complex immune dysregulation is often difficult. Given that our cohort studies show that elevation of IFN-α may precede diagnosis for decades, there is a need to develop a model which achieves stable elevation of IFN-α in the immune system of mice.

To address this gap, we created a new mouse model which allows IFN-α to be stably overexpressed from immune cells (Figure 4A, S4). IFN-α is elevated in both blood and tissues of SjD patients, and CD8^+^dendritic cells are an important source of IFN-α ^33^. Therefore we used a genetic strategy to overexpress Ifna4 in cells expressing the *Clec9a* gene that is a marker of conventional type I dendritic cells. We created a new mouse model “*R26mIfn*α*4*” which, upon Cre-mediated deletion of *LoxP-STOP-LoxP* cassette, expresses the early mouse Ifnα4 and EGFP (Figure 4A, S4). To force Ifnα4 expression predominantely in conventional dendritic cells type 1 in this model, we crossed *R26mIfn*α*4* mice with *Clec9a-Cre* mice ^27^, leading to an EGFP signal that was confined to a splenic population that is highly enriched for conventional type 1 dendritic cells (cDC1s) (Figure 4B). This resulted in high levels of Ifnα bioactivity and type I IFN signatures measured indirectly in peripheral blood, serum, and immune organs (Figure 4C, 4E S5), which caused immune disease in all *Clec9a^KI/WT^R26Ifn*α*4^KI/WT^* mice (from here on “KI” mice) (Figure 4D, 4E, S6A).

**Figure 4.**
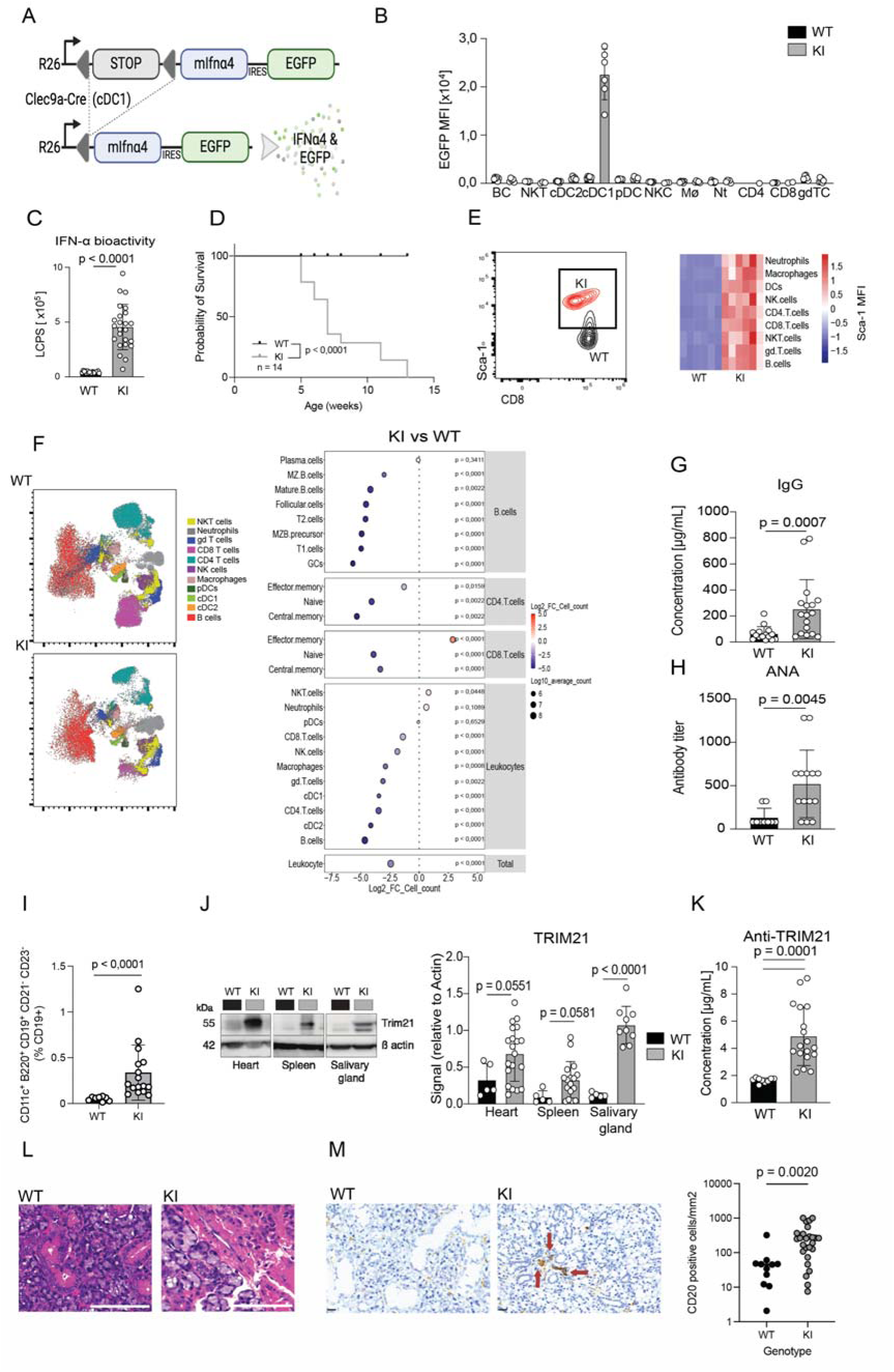
Constitutive IFN-α expression from conventional dendritic cell type 1 can initiate and recapitulate key features of the IFN-α Sjögren Disease endotype. A) Overview of the *Clec9a.cre^KI/WT^ x R26IFNa4^KI/WT^* mouse model. B) *Clec9a.cre^KI/WT^ x R26IFNa4^KI/WT(WT/WT)^* mice were sacrificed at 6 weeks of age. EGFP levels in the indicated spleen cell types shown as median fluorescence intensity (MFI) in different cell types (n=5/group). C) IFN-bioactivity in sera WT (n=17) and KI (n=24) mice. D) Kaplan Meier curve of mouse survival, n=14/group. E) Frequency of Sca-1^+^ CD8^+^ T cells in spleens of one WT vs. one KI mouse (upper) and a heatmap of Sca-1 MFI in the major splenic immune cell populations (down, n=6/group). F) High dimensional flow cytometry of KI (n=16) vs WT (n=8) spleens shown as UMAP plots (left) and as Log2 fold change (FC) of absolute counts for the depicted immune phenotypes in spleen cells (right). G) Concentrations of total IgG in sera of WT (n=15) and KI (n=17) mice were quantified by ELISA and increased in KI mice. H) Titers of IgG antinuclear antibodies (ANA) in sera of WT (n=9) and KI (n=14) mice were determined by indirect immunofluorescence, and were increased in KI mice. I)) Frequency of atypical memory B-cells (CD11c^+^B220^+^CD19^hi^CD21^-^CD23^-^), determined as a percentage of CD19+ cells, is increased in spleens of WT and KI mice. J) Representative western blot images (left) and quantification (right) of TRIM21 protein levels in heart, spleen and salivary glands of WT (n=5) and KI (n=9) mice. Black = WT, Grey = KI K) Concentration of TRIM21(Ro52) autoantibodies in sera of WT (n=9) and KI (n=18) mice were quantified by ELISA, and found to be increased in KI mice compared to WT mice. L) Chronic IFN-α4 elevation is associated with chronic inflammation and fibrosis of the salivary glands, the adjacent lymph nodes and periglandular fat. Salivary glands with surrounding adipose tissue were dissected from 53 mice (WT n=15, KI n=38), fixed in formalin, sectioned at 5uM, stained with haematoxylin and eosin, and examined by a consultant head and neck pathologist. Adjacent lymph nodes were included in 28 of the dissected salivary glands (WT n=7, KI n=21) and were included in the evaluation where available. Representative images of normal salivary ducts from a WT mouse and periductal fibrosis in a KI mouse are shown. Larger images and quantification in Supplemental Figure 7. Scale bar = 100um. M) Salivary glands in Clec9aR26Ifna4 mice (right) have significantly more CD20 positive cells than salivary gland tissue from wild type controls (left) (253 cells/mm2 vs. 45 cells/mm2 p = 0.002). Data are presented as a mean ± SD. Statistical significance was determined by Unpaired t-test or Mann-Whitney depending on the data distribution. The difference in the survival curve was determined by the log-rank test.

### Forced chronic expression of IFN-***α*** from dendritic cells recapitulates key features of the Sjögren disease IFN-***α*** endotype

Peripheral blood cytopenias including leucopenia, lymphopenia, and neutropenia were a key feature of the immune endotype found in people with SjD and elevated IFN-α. Therefore we used transcriptomic and flow cytometry to characterise lymphocyte populations in the mouse model. Among the top downregulated genes we found key B lymphocyte-regulating genes like Pax5 and Fcer2a (Figure S6B). Pseudo single-cell deconvolution of the bulk data using the quanTiseq pipeline ^34^ detected a strong reduction of lymphocyte signatures in spleens of KI mice compared to controls, suggesting that the downregulation of these cell-type specific transcripts was caused by a reduced abundance of the respective cell types (Figure S6C). To better understand the dynamics within the different splenic immune cell populations upon chronic exposure to cDC1-derived IFN-α4, we performed deep immuno-profiling of spleens using spectral flow cytometry. Indeed, KI mice presented with a pronounced leucopenia across the majority of immune cell types, most dominantly in B cells (Figure 4F), with evidence of Daxx-mediated block in B-cell development (Figure S7). In contrast to all other major leucocytes populations, numbers of CD8^+^CD44^+^CD62L^-^ effector memory T cells were substantially increased in KI vs WT spleens (Figure 4F), although depletion of CD8β cells did not significantly impact survival (Figure S8). Taken together with data from the SjD cohort, our analyses demonstrate that chronic systemic elevation of IFN-α contributes to the development of cytopenias, in particular lymphopenia.

Given our findings of an association between serum IFN-α concentrations and hypergammaglobulinaemia in SjD patients, we next examined immunoglobulin and autoreactive antibody formation in our mice and observed approximately two-fold higher IgG levels and higher ANA titers in KI in comparison with WT mice (Figure 4G & 4H). Thus, our data demonstrate that leukopenia, hypergammaglobulinaemia and ANA can be induced by chronic elevation of IFN-α. This is particularly notable given the profound lymphopenia in KI mice, including strong suppression of B-cell counts, suggesting a preferential survival and activation of autoreactive B cells in the presence of chronic IFN-α. Consistent with this hypothesis, we identified a significant increase in the proportion of atypical memory B-cells (B220+CD21-CD23-CD11c+CD19hi) in KI mice (Figure 4I).

Informed by the particularly strong association we observed between IFN-α and TRIM21 autoantibodies in people with SjD we looked for evidence of TRIM21 upregulation in our mice. Indeed, the TRIM21 protein levels were increased in total cell lysates of spleens and hearts from KI compared to WT controls (Figure 4J). Consistently, we found increased concentrations of TRIM21 autoantibodies in the serum of KI, but not of WT mice (Figure 4K). The increased abundance of TRIM21 in tissues seems to underly the autoantibody production, as we failed to detect circulating TRIM21 antigen in the serum of KI mice (Figure S6D), similar to our observation in SjD patients (Figure 3E). We observed high *TRIM21* expression together with histological changes in the salivary glands (periductal fibrosis), and inflammatory change in the periglandular fat of KI mice, respectively (Figure 4L, 4M & S9). We observed a significant increase in B cell number within salivary glands (Figure 4M).

### Blockade of IFN-α signalling ameliorates specific aspects of the IFN-α endotype

Systemic blockade of IFN-α signalling has been approved for the treatment of type I interferon-associated autoimmune diseases such as SLE ^14^. To assess the therapeutic effect on Sjögren-associated immune alterations, we treated KI mice with the IFNAR1 blocking antibody MAR1-5A3 three times per week for up to twelve weeks (Figure 5A). We observed reduction in molecular markers of interferon signalling (Figure. 5B, 5C, S6A) and clinical improvement with longer survival (Figure 5D). Similarly, cytokine profiles, cytopenias and abnormal haematological parameters of the treated mice were all improved compared to untreated mice (Figure 5E, 5F, S6E). However, total IgG levels and ANA titers remained unaffected after four weeks and slightly increased after 12 weeks by MAR1-5A3 treatment compared with isotype controls (Figure 5G, 5H). Similarly, Trim21 antibody levels were not reduced after 4 weeks of MAR1-5A3 treatment (Supplementary Figure 10). Although we observed a reduction of the ISG signature (Figure S6A, S6G), transcript levels and intracellular protein abundance of TRIM21 did not revert to levels found in untreated wild-type mice through pharmacologic inhibition of IFNAR signalling (Figure S6F, Figure 5I). These results suggest that although important aspects of disease caused by life-long elevation of IFN-α can respond to interferon receptor blockade, some features can persist.

**Figure 5.**
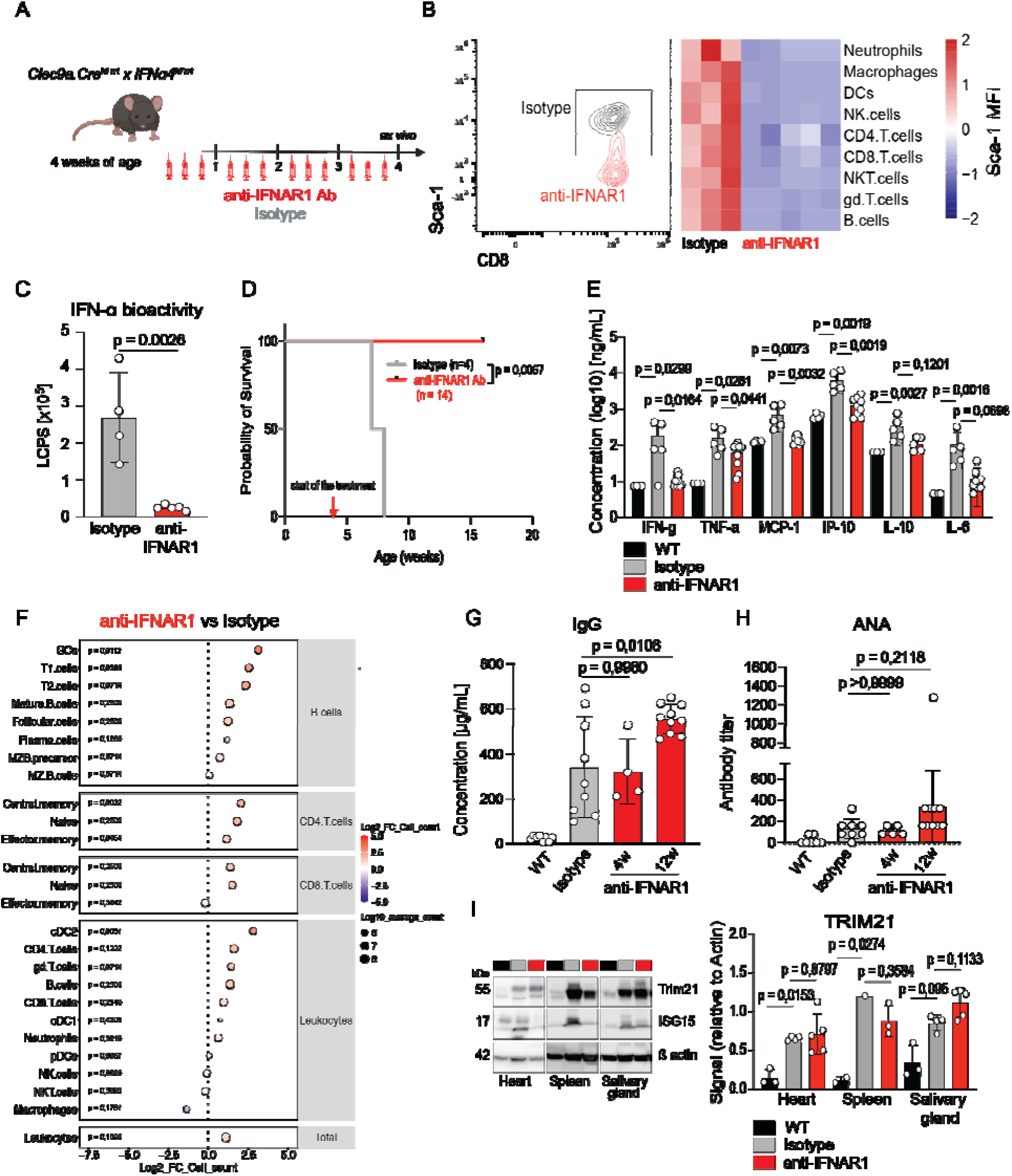
The chronic IFN-α driven endotype partially responds to pharmacologic inhibition of type I interferon signalling. (B) Experimental design with overview of treatment timings, treatment arms (anti-IFNAR1 Ab versus Isotype) and endpoint. (C) Contour plot of Sca-1^+^ CD8^+^ T cells in spleens of one Isotype vs. one anti-IFNAR1 treated *Clec9a.cre^KI/WT^ R26IFNa4^KI/WT^*(KI) mouse (left). MFI of Sca-1 among depicted splenic populations shown as a heatmap (right, Isotype n=3, anti-IFNAR1 n=6). (D) IFN bioactivity in sera from Isotype (n=4) and anti-IFNAR1 (n=5) treated mice. (E) Kaplan Meier survival curve, Isotype (n=4) and anti-IFNAR1 (n=14) treated mice (F) Cytokine levels in the sera of WT (n=4), Isotype (n=5) and anti-IFNAR1 (n=9) treated mice. (G) High dimensional flow cytometry of Isotype (n=3) and anti-IFNAR1 (n=5) treated KI mice shown as Log2 fold change (FC) of absolute counts for the depicted immune phenotypes in spleen cells. (H) Concentration of IgG in sera of WT (n=8), Isotype (n=9), 4 weeks (n=4) and 12 weeks (n=10) anti-IFNAR1 treated mice quantified by ELISA. (I) Titers of IgG antinuclear autoantibodies (ANA) in sera of WT (n=7), Isotype (n=8), 4 weeks (n=5) and 12 weeks (n=10) anti-IFNAR1 treated mice were determined by indirect immunofluorescence. (J) Representative western blot images (left) and quantification (right) of TRIM21 protein levels in heart, spleen and salivary glands of WT (n=2), Isotype (n=1) and anti-IFNAR1 (n=3) treated mice.

## DISCUSSION

Developing targeted treatments for SjD requires the identification of the dysregulated immune pathways which drive disease, together with an understanding of how activation of these pathways may vary between individuals. Here, we have adopted an interdisciplinary approach, combining emerging protein detection technologies with cohort studies and experimental modelling, to provide new insights into SjD heterogeneity, and show how decades-long elevation of IFN-α can drive an endotype of SjD.

Collectively, our combined Simoa-Olink studies in UKPSSR and UK Biobank cohorts show that IFN-α is elevated in a substantial subpopulation (∼60%) of individuals with SjD, and this elevation can precede clinical manifestations by over a decade. This chronic IFN-α elevation is associated with a distinct interferon-related immune endotype characterised by blood cytopenias, IgG production and autoantibody production. Elevated IFN-α is also associated with the development of autoantibodies commonly found in SjD, in particular antibodies directed against TRIM21. Such immune changes (notably reduced TRIM21) can be identified in the period prior to diagnosis in broad-capture proteomic datasets and have potential utility for the early identification of populations at-risk of developing SjD ^35^. Importantly, IFN-α concentrations are not associated with measures of disease activity in our UKPSSR cohort, nor with pattern of organ activity (as measured with ESSDAI domains). As such, SjD patients with elevated IFN-α concentrations are clinically similar to those with normal IFN-α concentrations, yet are immunologically distinct. This highlights the critical role for optimised biomarkers for accurate detection of those who could benefit from therapeutic approaches which target activation of the type I IFN response.

Our ultrasensitive IFN-α Simoa assay highlights that 40% of patients do not have elevated IFN-α, despite meeting the same diagnostic criteria, experiencing similar levels of disease activity with similar patterns of organ activity. The precision of our Simoa assay, specifically optimised to detect IFN-α at very low concentrations, offers confidence that IFN-α concentrations in this population are within the range of healthy controls, rather than simply below the limit of detection of the assay. Accurate identification of this subpopulation will be important in supporting discovery work to identify identify interferon-independent pathways that can initiate and drive SjD.

To explore the origins of the IFN-α endotype, we have developed and characterised an oligoprotein interferon signature approach using Olink proteomics which allows us to identify elevations of IFN-α well over a decade prior to SjD diagnosis in the UKBiobank-PPP. Our use of the oligoprotein score, which we term SIRO, to trace the origins of IFN-α highlights how early identification of at-risk individuals could identify a therapeutic window which could be targeted for disease prevention. Our SIRO score has a number of potential uses, in particular in the interrogation of large population-based biobanks where measurement of IFN-α has traditionally been challenging because of the need for high volumes of plasma. Like all interferon signatures it is not a perfect surrogate but has the potential for further refinement, including use with other autoimmune diseases and in understanding how IFN signatures can vary between different IFN-associated autoimmune diseases. Our detection of oligoprotein interferon signatures up to 14 years prior to SjD diagnosis highlights that activation of the type I interferon response can occur for decades before a final diagnosis is made, highlighting the need to understand the downstream immunological consequences of persistent immune dysregulation.

While cohort studies can provide evidence of association between specific cytokines and immunological phenotypes, causality is notoriously difficult to prove. Biological models can play an important role in demonstrating biological plausibility, and provide insights into human epidemiological studies when a causal role for a cytokine in driving aspects of disease is being considered. To address this gap in knowledge, we developed a novel transgenic mouse model of chronic Ifn`U`4 overexpression. Our model provides proof of principle that these immunological features can be both initiated and driven by chronic IFN-α production - and can also be therapeutically targeted through IFNAR blockade. This *in vivo* approach represents a significant advance in studying the immunological sequelae of chronic IFN-α overproduction, modelling the decades-long stable elevation of IFN-α we observed in people with SjD. While the cellular source of IFN-α in patients with autoimmunity is complex ^12^, overexpression of *Ifn*α*4* in Clec9a-expressing cDC1 cells achieves an increase in tissue IFN-α, as well as elevated blood IFN-α.

From the perspective of addressing unambiguously the downstream consequences of very high levels of IFN-α in the peripheral immune system, this model offers considerable advantages over established SjD mouse models. Importantly we show that near-lifelong elevation of IFN-α causes and can drive key features of the IFN-α endotype we have identified (e.g. cytopenia, hypergammaglobulinaemia, autoantibody generation). This includes the generation of autoreactivity directed against the key SjD autoantigen TRIM21. In our mouse model we observed elevation of TRIM21 in salivary glands, as well as other tissues. Highlighting the disease-relevance of this observation, elevation of TRIM21 protein has also been observed in human SjD salivary glands, and this elevation is associated with the level of inflammation ^36^.

As well as providing evidence to support a role for IFN-α in driving SjD, our mouse model also provides mechanistic insights into aspects of the IFN-α endotype. In particular, our prediagnostic studies highlight the need to understand the consequences of longterm IFN-α elevation and our model fills an important gap in our tools to study this. For example, our mouse studies illustrate the profound influence of chronically elevated IFN-α on B-cells, where we observed IFN-induced block in B-cell development possibly via Daxx, as well as an increase in proportions of atypical memory B-cells and tissue-resident B-cells. Further detailed mechanistic studies of this IFN-α B-cell dysregulation, and the degree to which these persist despite IFNAR blockade, will be of considerable interest. Our transgenic mouse findings also demonstrate that the immunological consequences of chronic IFN-α elevation can be challenging to reverse through receptor blockade. While therapeutic blockade of IFNAR signalling offers some efficacy, certain aspects of the immune dysregulation (e.g. autoantibody formation) cannot be rapidly reversed with receptor blockade alone. Such findings may help explain some of the mixed results of IFNAR blockade in interferon-associated autoimmune diseases such as SLE^13,14^ and may help understand which aspects of interferon-driven immune disease might be predicted to respond clinically to receptor blockade, and which ones may not. This further highlights a potential need for early diagnosis and intervention, which is facilitated by our finding that broad capture proteomics can be used for early identification of individuals at risk of developing SjD. The significant improvement in survival in the mice treated with IFNAR blockade contrast with the persistence of aspects of the autoantibody response, which suggest that these autoreactive antibodies may not be mediating important disease-specific aspects in this model. More generally these results show how B-cell lineage dysregulation has the potential to persist in the scenario of near lifelong IFN elevation, even if IFNAR signalling is blocked. Integration of these biological principles with existing and ongoing trials of IFNAR blockade in systemic autoimmunity will be of substantial interest.

Our combined human and mouse model studies highlight that an important feature of the IFN-α endotype is the development of autoreactivity against the Ro52 antigen, TRIM21. TRIM21 is considered to be a classic autoantigen for SjD as well as other autoimmune conditions including SLE. Notably genetic deletion of TRIM21 leads to systemic autoimmunity ^37^. The dysregulated dynamics observed here in SjD are of particular interest and suggest that IFN-α may play a key role in the initiation of the perturbed state where there is high intracellular TRIM21 expression yet low serum levels that may be related to TRIM21 antibodies. The role of reduced TRIM21 in the initiation and propagation of autoimmunity represents an important area for further study.

There is a well-described overlap between SjD and other genetic and complex immune-mediated diseases such as monogenic interferonopathies and SLE, where elevated IFN-α is observed in an even higher proportion of affected individuals (Rodero et al. 2017). Our findings are also relevant to these disorders and highlight that identifying SjD associated IFN-α driven changes in SLE, such as dysregulated TRIM21 dynamics, can predict the development of co-morbid SjD. Our mouse model aims to identify the downstream consequences of elevated IFN-α rather than develop a SjD model *per se*, and some of the findings may be of relevance to IFN-α driven immunopathology in other autoimmune disease such as SLE.

Our interdisciplinary study has a number of limitations. A limitation of our human cohort work is that the UKBiobank and UKPSSR are distinct studies. While we have prediagnostic serum analyses available for UKBiobank, there are no prediagnostic UKPSSR samples available. However the association between IFN-α and SjD is replicated in both independent studies which strengthens our findings. Because of limitations on testing large biobanked cohorts (sample volume for Simoa is ∼100ul, which is a current limiting factor for testing in biobanks with depletable serum resources) we have used an indirect measure of IFN-α using an oligoprotein signature derived from broad capture proteomics, which is a reasonable but imperfect proxy for IFN-α and can be analysed in >45000 UK-PPP participants. There are also limitations on the statistical power of our UKPSSR cohort (n=177) and therefore future studies in large cohorts may be able to identify subtle clinical associations which we have not been able to detect.

Finally, there are multiple IFN-α subtypes, all of which act on a common receptor IFNAR. Our Simoa assay is a multiple subtype assay, which correlates well with ISG activation downstream of IFNAR, but does not give details of which subtypes are elevated, Taken together our findings have implications for precision medicine and stratified medicine approaches in SjD. Firstly, through our interdisciplinary approach to establishing IFN-α as a driver of an SjD endotype, we have identified IFN-α as an important potential therapeutic target. Secondly we have highlighted that targeting IFNAR through therapeutic blockade can lead to striking improvements in some aspects of disease, yet others may be unaffected. This is likely due to the establishment of interferon-independent aspects of autoimmunity and should be carefully borne in mind when evaluating the results of trials which target the receptor. Notably IFNAR blockade in SLE led to mixed phase 3 results, and similar studies in SjD are in progress. A detailed biological understanding of how specific components of the IFNAR-driven endotype arise and respond to treatment could help optimise trial design. Thirdly, our work develops further a number of practical biomarkers which can be employed in clinical research and practice in order to stratify SjD and other IFN-α associated immune diseases along precision medicine principles. As well as Simoa with improved sensitivity, we have also shown how an Sjögren-relevant interferon score can be developed from broad capture proteomics datasets, opening up large healthcare datasets such as UK Biobank to the study of type I IFN biology, which has been difficult to achieve to date. Finally we have identified a potential need for early diagnosis and intervention. Our deconvolution of prediagnostic proteomic datasets have shown that Olink derived IFN-related markers can predict onset of SjD, including “secondary SjD” in patients with SLE – and our experimental studies have highlighted how longstanding (and potentially undiagnosed) elevated IFN-α levels can lead to the establishment of persistent immune dysregulation that can respond only partially to receptor blockade. As such, these findings have the potential to support “precision prevention” approaches in SjD, which enable identification of at-risk individuals early in their disease course who may benefit the most from targeted treatment. Therefore our work develops tools which are of practical use for disease stratification, trial design and risk prediction.

In summary our results identify an IFN-α endotype of SjD, driven by elevations of IFN-α that can persist for decades. Drugs targeting this pathway in SjD should be prioritised for interventional trials, which should be designed to account for the immunological heterogeneity of this condition.

## Acknowledgments

We thank Thomas Wunderlich for the pSERC vector, the Transgenic Core Facility of the MPI-CBG Dresden for electroporation of ES cells as well as the blastocyst injections, Flow Cytometry Core Facility at the Institute of Experimental Immunology, University Hospital Bonn, and the Next Generation Sequencing Core Facility of the Medical Faculty/West German Genome Center Bonn at the University of Bonn for providing support and instrumentation for the 3’mRNASeq. We thank Barbara Uteß, Christina Hiller and Livia Schulze for excellent technical assistance. FN has undertaken clinical trials and provided consultancy or expert advice in the area of Sjogren disease to the following companies: GlaxoSmithKline, MedImmune, UCB, Abbvie, Takeda, Resolves Therapeutics, Sanofi, Novartis, Bain Capitals and Argenx. All other authors have nothing to declare.

---

**Table 1.** Demographics and clinical characteristics of the UK Primary Sjögren Syndrome Registry (UKPSSR) cohort.

| **Demographics** | **UKPSSR IFN-α Simoa cohort (n=177)** |
| --- | --- |
| **Age (median, (IQR))** | 57.5, (46-65) |
| **Female sex, n (%)** | 163/177 (92%) |
| **BMI (median, (IQR))** | 25.9, (23.0, 29.1) |
| **SjD duration in years (median, (IQR))** | 4, (2, 8) |
| **Ethnic background white %** | 162/177 (91%) |
| **Ethnic background non white %** | 15/177 (9%) |
| **Baseline Disease activity ESSDAI (median, (IQR))** | 3, (1,6) |
| **Baseline symptom score ESSPRI (median, (IQR))** | 5.5, (2.33, 7.33) |

