## Supplemental data for "Decades-long elevation of interferon-α drives a Sjögren disease endotype: an interdisciplinary study"

* and ** denote equal contributions

**SUPPLEMENTARY METHODS**

**Whole blood transcriptomics.**

Whole blood was collected in PAXgene tubes (Becton, Dickinson and Company, Oxford) at the time of recruitment to the UKPSSR and stored at -80°C until RNA extraction and then assayed with the Illumina Human HT-12 v4 Expression BeadChip (Illumina, Cambridge, United Kingdom). Globin signal suppression was performed using the Affymetrix Globin reduction protocol (Affymetrix, Santa Clara California, United States of America). Analysis was performed using Bioconductor libraries in the R environment for statistical computing.

**Chaussabel module scores**

The Chaussabel method was used to calculate transcriptomic activity scores (Chaussabel et al., 2008). Chaussabel modules represent groups of genes that are clustered according to the physiological and immunological state of 965 human leukocyte transcriptomes. Three Chaussabel modules (M1.2, M3.4 and M5.12) are typically considered to reflect ISG expression (Chaussabel et al., 2008; Tarn et al., 2019). For each Chaussabel module, an average expression score was calculated using pooled data from all 177 SjD patients. Each patient was assigned a score relative to this average, used for the correlation with blood IFN-α.

**Future MS**

Serum samples were sourced from a further eight healthy controls were sourced from the Future MS study, a national Scottish observational cohort study. Participants gave informed written consent at time of recruitment. Venepuncture was performed on the same day and serum collected using ETDA tubes, all samples were stored in Eppendorf tubes at -80°C until the time of analysis. Research ethics approval was granted by the NHS South East Scotland Research Ethics Committee [02]: [REC 15//SS/0233]).

**Definition of cases in the UK Biobank**

A case of SjD was defined as at least one diagnostic code from inpatient hospital, primary care or death records, or any self-reported diagnosis at any assessment visit. First occurrence fields in UKB, which map all available diagnostic sources (i.e., self-reports at baseline and follow-ups, hospital admissions, death registries and primary care) to International Classification of Diseases (ICD) v10 3-digit diagnosis (biobank.ndph.ox.ac.uk/showcase), were used for multiple sclerosis (ICD v10 G35 [multiple sclerosis]; fields 131042, 131043) and haemorrhoids (ICD v10 I84 [haemorrhoids] and K64 [haemorrhoids and perianal venous thrombosis]; fields 131404, 131405, 131650, 131651). Health-related outcomes were assessed using the June 2023 data release.

| **System** | **Code** | **Definition** |
| --- | --- | --- |
| ICD v10 | M35.0 | Sicca syndrome [Sjögren] |
| ICD v9 | 710.2 | Sicca syndrome |
| Read v2 | F3967 | Myopathy due to Sjögren disease |
| Read v2 | H57y3 | Lung disease with Sjögren disease |
| Read v2 | N002. | Sicca (Sjögren) syndrome |
| Read v3 | H57y3 | Lung disease with Sjögren disease |
| Read v3 | N002. | Sicca (Sjögren) syndrome |
| Read v3 | X705A | Primary Sjögren syndrome |
| Read v3 | X705B | Primary Sjögren syndrome with organ/system involvement |
| Read v3 | X705C | Primary Sjögren syndrome with multisystem involvement |
| Read v3 | X705D | Secondary Sjögren syndrome |
| Read v3 | X705E | Secondary Sjögren syndrome with organ/system involvement |
| Read v3 | X705F | Secondary Sjögren syndrome with multisystem involvement |
| Read v3 | XE1Ge | Sjögren disease |
| Self-report | 1382 | Sjögren syndrome/sicca syndrome |

**HLA association with Olink proteomics in the UK Biobank**

We explored whether the previously reported association of HLA-DQA1*05:01 with SjD (Lessard et al., 2013) and interferon-alpha levels in SjD (Trutschel et al., 2022) could be replicated in UKB using oligoprotein interferon signatures. HLA-DQA1*05:01 allele dosage was defined from genotype array data using imputed HLA values generated with HLA*IMP2 (UKB field 22182). HLA associations with SjD and SIRO (z-scored) were respectively assessed using logistic regressions with Firth penalization and linear regressions, both adjusted for age, sex and the 10 first genetic principal components.

**Indirect immunofluorescence for the detection of ANAs**

Mouse 3T3 fibroblasts were seeded at a density of 8 × 10^4^ cells/mL on glass slides (Carl Roth) and grown overnight in DMEM followed by fixation and permeabilization with ice-cold 90% methanol / 10% acetone for 60 min at -20°C. Slides were incubated for 1h at room temperature with serial dilutions of sera from 1:80 - 1:1280. After washing, bound antinuclear antibodies (ANA) were detected using a polyclonal goat anti-mouse IgG-Alexa Fluor 488 (Poly4053, 1:500, Biolegend) for 1h at RT. Lastly, slides were stained with DAPI and mounted with ProLong Gold Antifade Mountant (Thermo Fisher). Immunofluorescence patterns were analyzed on a Thunder microscope (Leica) and were assessed by two independent researchers.

**Flow cytometry**

Homogenized spleen cell suspension was subjected to erythrolysis and passed through a 70-μm cell filter. Following washing with ice-cold FACS buffer (PBS/1%FCS/ 2mM EDTA), cells were incubated with anti-CD16/CD32 (1:50, Biolegend) for 10 min to block Fc receptors and then stained with antibody mixture in FACS buffer at 4°C for 30 minutes. After two washing steps, cells were processed on spectral cell analyser ID7000 5-Laser (Sony Biotechnology). Data were analyzed by FlowJo software (BD Biosciences) whereas the figures are made in R studio (version 4.2.2). UMAPs were created in FlowJo software (BD Biosciences) by using 8 samples per group, concatenated to 700 000 event per sample (for WT/KI comparison), whereas for the treatment, 3 samples per group were concatenated and normalized to 1 million events per sample. For the cell count of WBC in the whole blood, samples were erythrolysed and stained with anti-CD45 APC-Cy7 antibody for 30min followed by two washing steps and resuspension in DAPI. Spleen cell counts were recorded after DAPI staining on MacQuant 10 analyser (Miltenyi).

Antibodies used for spectral analysis and cell counts are summarized in the table below.

| **Antibody** | **Clone** | **Dilution** | **Company** |
| --- | --- | --- | --- |
| CD45R/B220 NovaFluor™ Yellow 660 | RA3-6B2 | 1:100 | Thermo Fisher |
| CD45R/B220 BV 711 | RA3-6B2 | 1:50 | Biolegend |
| Ly-6A/E (Sca-1)  Spark NIR 685 | D7 | 1:100 | Biolegend |
| I-A/I-E Spark Violet™ 538 | M5/114.15.2 | 1:50 | Biolegend |
| Ly-6G/Ly-6C (Gr-1)  BV 570 | RB6-8C5 | 1:100 | Biolegend |
| CD45 Alexa Fluor 700 | 30-F11 | 1:300 | Biolegend |
| CD45 APC-Cy7 | 30-F11 | 1:300 | BD |
| CD19 BV 650 | 6D5 | 1:300 | Biolegend |
| CD64 (FcγRI) BV 605 | X54-5/7.1 | 1:20 | Biolegend |
| CD138 (Syndecan-1)  PerCP/Cy5.5 | 281-2 | 1:100 | Biolegend |
| CD172a (SIRPα) PE/Dazzle 594 | P84 | 1:50 | Biolegend |
| GL7 APC | GL7 | 1:20 | Biolegend |
| XCR1  BV785 | ZET | 1:100 | Biolegend |
| CD95 (Fas) PE/Cy7 | SA367H8 | 1:20 | Biolegend |
| NK-1.1 PE/Fire 700 | S17016D | 1:100 | Biolegend |
| CD62L PE/Fire 810 | W18021D | 1:100 | Biolegend |
| IgM BV421 | RMM-1 | 1:50 | Biolegend |
| IgD PE | 11-26c.2a | 1:100 | Biolegend |
| CD3  Spark Blue 550 | 17A2 | 1:100 | Biolegend |
| CD8a  Spark UV 387 | 53-6.7 | 1:50 | Biolegend |
| CD25 PE/Fire 640 | PC61 | 1:100 | Biolegend |
| CD44 Spark YG 593 | IM7 | 1:50 | Biolegend |
| CD11b Pacific Blue | M1/70 | 1:100 | Biolegend |
| γδ TCR BUV661 | GL3 | 1:50 | BD |
| Siglec-H BV480 | 440c | 1:100 | BD |
| CD21/CD35 BUV615 | 7G6 | 1:50 | BD |
| CD11c BUV563 | N418 | 1:50 | BD |
| CD23 BUV805 | B3B4 | 1:100 | BD |
| CD43 BUV737 | S7 | 1:50 | BD |
| CD4 BUV496 | GK1.5 | 1:50 | BD |
| Zombie NIR^TM^ Fixable Viability Kit |  | 1:1000 | Biolegend |

Defining the cell populations:

Spleen B cells are defined according to the references (Morgan D. et al. 2022, Peschke K. et al. 2016, Shinall S.M. et al., 2000**)** as follow:

*Mature* (CD19^+^ B220^high^ IgD^high^), *T1* (CD19^+^ B220^high^ IgM^high^ IgD^low^ CD21^low^), *T2* (CD19^+^ B220^high^ IgM^high^ IgD^high^ CD21^low^), *MZB precursor* (CD19^+^ B220^high^ IgM^high^ IgD^high^ CD21^high^), *MZB* (CD19^+^ B220^high^ IgM^high^ IgD^low^ CD21^high^ CD23^low^), *Fo* *B* (CD19^+^ B220^high^ IgM^low^ IgD^high^ CD21^int^ CD23^high^), *GC* (CD19^+^B220^high^ CD95^+^ GL7^+^), *Plasma cells* (CD19^+^ B220^low^ IgD^low^ CD138^+^).

Dendritic cells are defined as follow:

*pDCs* (CD11c^+^ MHCII^+^ Siglec H^+^), *cDC1* (CD11c^+^ MHCII^+^ XCR1^+^ CD8a^+^), *cDC2* (CD11c^+^ MHCII^+^ Sirpa^+^ CD4^+^).

*T cells were divided depending on the expression of CD44 and CD62L into: Effector memory T cells* (CD44^+^ CD62L^-^), *Central memory T cells* (CD44^+^ CD62L^+^), *Naive T cells* (CD44^-^ CD62L^+^).

**Western blot**

Whole spleen, salivary gland and heart were lysed in RIPA buffer containing Phenylmethanesulfonyl fluoride (PMSF) (Carl Roth), cOmplete EDTA-free protease inhibitor cocktail (Sigma Aldrich), and PhosStop (Sigma Aldrich). The concentration of proteins was measured by Bradford assay and 50 ug of total protein was separated by SDS-PAGE followed by semi-dry transfer to a PVDF membrane. After blocking in 5% milk/ TBS-T solution, membranes were incubated with a monoclonal mouse anti-mouse Trim21 antibody (1:2000, Proteintech), monoclonal mouse anti-mouse ISG15 (1:1000, Santa Cruz Biotechnology)  and rabbit anti-mouse β-actin (1:5000, Proteintech) overnight at 4°C or mouse anti-mouse β-actin-HRP antibody (1:4000, Santa Cruz Biotechnology) for 1h at RT. Detection was performed using HRP-conjugated secondary horse anti-mouse IgG antibodies (1:5000, Cell Signaling) followed by addition of ECL blotting substrate (Thermo Fisher Scientific). Images were recorded on an Odissey Fc (Li-Cor) device.

**Serology: ELISA, IFN type I bioassay, Cytokine bead array (CBA) assay**

Sera of mice were collected at the age of 6 weeks and analysed for the presence of anti-SSA antibodies (Signosis), total IgG (Thermo Fisher) and total Trim21 protein (Hölzel) by ELISA according to the manufacture’s protocols. IFN type I bioactivity was quantified by overnight incubation of sera (1:4 dilution in DMEM) with LL171 cells (Uzé et al., 1994) that stably express an ISRE-dependent-luciferase. LL171 cells were lysed in the Cell Culture Lysis Reagent (Promega) for 20 min and luciferase activity was measured on Multimode Microplate Reader (Promega) after the addition of D-Luciferin (Sigma) substrate. For the measurement of cytokine levels sera were collected after 4 weeks of anti-IFNAR1 (and Isotype) treatment and were analysed by LEGENDplex Mouse Anti-Virus Response Panel (13-plex) kit (Biolegend) according to the manufacturer’s protocol. Samples were recorded on Attune NxT flow cytometer (Thermo Fisher) and quantification was done on the LegendPlex platform recommended by the manufacturer.

**Haematoxylin and eosin (H&E) and CD20 immunohistochemical quantification of salivary glands**

Salivary glands with surrounding adipose tissue were dissected from 53 mice (WT n=15, KI n=38), fixed in formalin, sectioned at 5uM, stained with *H&E* and examined by a consultant head and neck pathologist. Adjacent lymph nodes were included in 28 of the dissected salivary glands (WT n=7, KI n=21) and were included in the evaluation where available.

For CD20 quantification, salivary glands and their adjacent lymph nodes were dissected and fixed in formalin then embedded in paraffin wax. Sections of 5uM were cut and mounted onto slides with anonymised IDs for staining and analysis. The CD20 stain was performed with Abcam ab64088 at a 1:250 concentration using a Leica Bond RX autostainer instrument.

Quantification of the CD20 stain was performed with QuPath open access image analysis software version 0.5.1-x64. The salivary gland, lymph node, periglandular fat, and interstitium were defined with regions of interest and then the QuPath positive cell detection function was used to identify positive cells based on the optical density of DAB surrounding the haematoxylin stained nuclei with the following settings, Background nucleus size 8uM, median filter radius 0uM, sigma 1.5uM, minimum area 10uM2, maximum area 400uM2, intensity threshold 1, max background intensity 2, cell expansion 5uM to include the nucleus, threshold 1 = 0.2, threshold 2 = 0.4, threshold 3 = 0.6. The number of positive cell/mm2 were exported from QuPath to a csv file. Where an individual salivary gland required multiple ROIs to capture all salivary gland tissue, the mean cells/mm2 across all ROIs was calculated and used for analysis. The KI (n=26) and WT (n=11) groups were compared with a Mann Whitney U Test in Graphpad Prism v10.

**Bulk RNA sequencing**

Total RNA was isolated from the whole spleen with Trizol according to manufacturer’s protocol (Thermo Fisher) followed by DNAse treatment.  cDNA libraries were prepared at the Next Generation Sequencing Core Facility of the Medical Faculty/West German Genome Centre Bonn at the University of Bonn using the QuantSeq 3´-mRNA Library protocol from Lexogen. Sequencing was performed on an Illumina NovaSeq 6000 platform to generate 1x100bp reads resulting in an average depth of 10 million Raw Reads per sample. Count files (fastq) were pre-processed for quality and adapter trimming using cutadapt (v.4.9) (Martin M., 2011)**.** High quality reads were aligned using Salmon (v1.7) (Patro et al., 2017) to the mm10 transcriptome (GCA_000001635.2) and checked for quality using Fastqc (v.0.12.0) and Multiqc (v1.11). Differential gene expression analysis was carried out using DESeq2 (v.1.34.00) (Love et al, 2017). Biological outliers (1 WT and 2 KI samples) were determined using PCA and heatmap visualization and removed for final analysis. Two sequencing runs were conducted, and raw data was batch-corrected using CombatSeq in sva package (v.3.38.0) (Zhang, Y., 2020). Comparisons were run using LRT  test and multiple testing correction was implemented using the Benjamini Hochberg method. The cutoff for differentially expressed genes (DEGs) was *Padj* < 0.05. Heatmaps were visualized using the pheatmap package (v.1.0.12) (For detailed steps see ‘Code availability’).

**Cell type deconvolution**

Relative enrichment of immune cell types in whole-spleen RNAseq samples was determined using DESeq2 normalized expression values. Gene names checked against the *base* database with standard parameters of the Immunodeconv package (Sturm et al., 2020).

(See ‘Code availability’)

**Data availability**

Raw and processed RNAseq data that support the findings of this study have been deposited into Gene Expression Omnibus (GEO) accession numbers GSE273095 and GSE273098

**Code availability**

Code for analysis and generating figures is available at: https://github.com/The-Behrendt-Lab/R26mIfna4_mouse

**Statistical analysis**

Statistical analysis and graphing of the single-molecule ELISA and clinical data were performed on graphpad prism software v10.0. Spearman’s Test was used to assess correlation between tests. To assess significance between two variables, the non-parametric Mann-Whitney U test was employed with the threshold for a significant result set at p<0.05.

In mouse studies normal distribution was assessed by Shapiro-Wilk normality test and comparison between variables is done by unpaired Student’s t-test or Mann-Whitney non-parametric test for comparing two variables, whereas the comparison of multiple variables is done by one-way ANOVA or Kruskal-Wallis test depending on the distribution of data. In flow cytometry data, significant outliers, detected by GraphPad Prism outlier calculator, were excluded from the analysis. P < 0.05 was used as the cut-off for significance in all tests, as follows: *, P < 0.05; **, P < 0.01; ***, P < 0.001; ****, P < 0.0001. All error bars represent SD.

**SUPPLEMENTARY FIGURES**

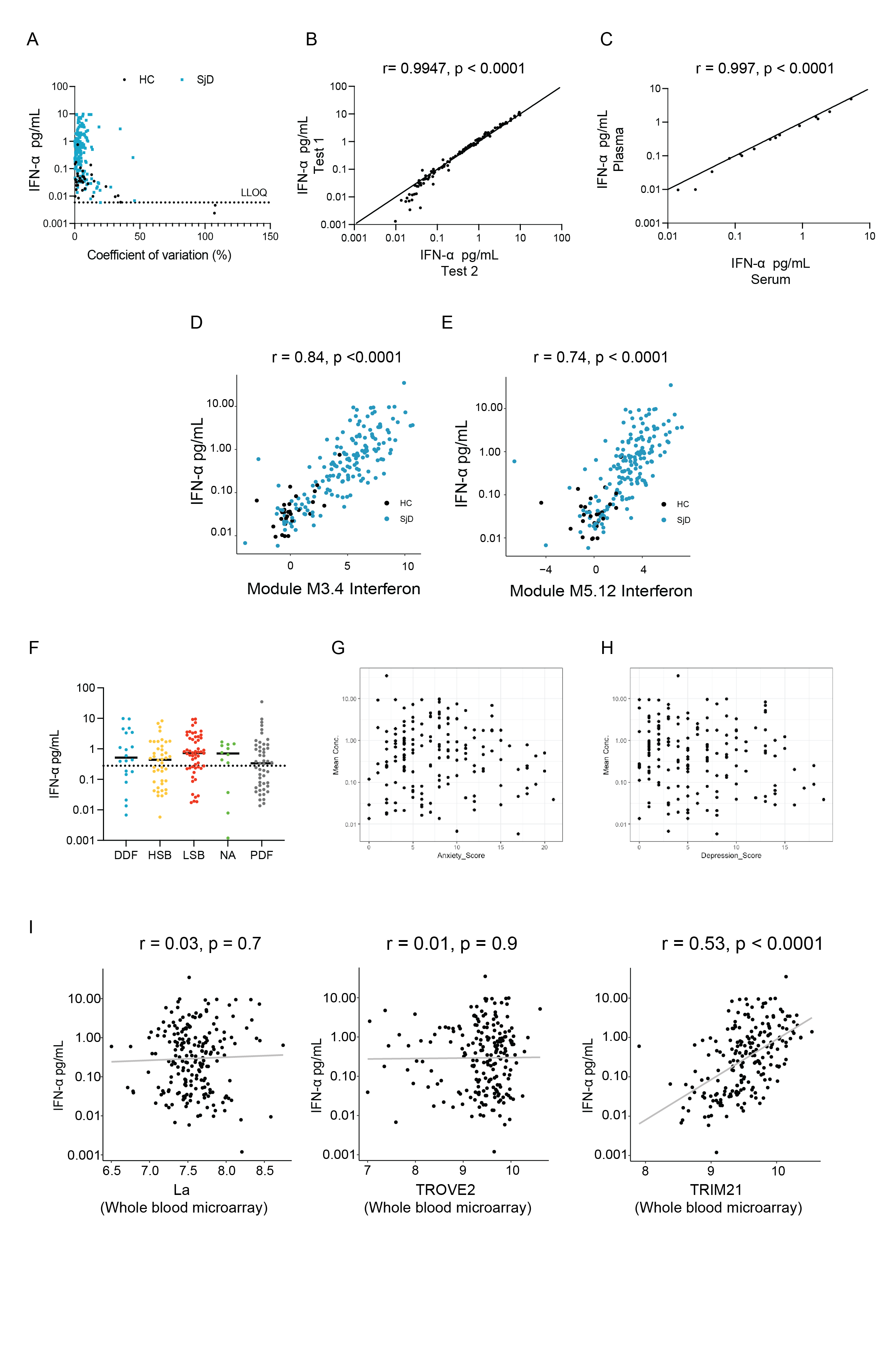

**Supplemental Figure 1. Single molecule array digital ELISA precisely and reliably measures the concentrations of IFN-α in human serum.**

(A) The pan IFN-α Simoa has excellent intra-assay reproducibility and a lower limit of quantification in the femtomolar range. Coefficients of variation were calculated for each serum sample's IFN-α across two replicates. The LLOQ was calculated using the concentration at which the coefficient of variation was < 20% for each batch, the mean LLOQ across all tests was 0.0058 pg/ml.

(B) The pan IFN-α Simoa has excellent inter-assay reproducibility. Paired samples from the same venepuncture draw were stored and prepared using identical protocols and subject to quantification of IFN-α in two separate tests. There was a strong correlation between test 1 and test 2 (Spearman's rank r = 0.9948, p < 0.0001)

(C) Correlation of IFN-α protein measured by Simoa in paired plasma and serum samples from 15 volunteers (Spearman's rank r = 0.997, p < 0.0001). These samples were collected under approval granted from the NHS East of Scotland. Research Ethics committee (REC01) as part of the Scottish Regenerative Neurology Tissue Bank Project (Reference: 15/ES/0094, 15/SS/0233 & 15/ES/0094). Participants gave informed consent to participate in the study before taking part.

(D and E) A microarray for the Chaussabel modules M3.4 and M5.12 was used to quantify the transcriptomic response to IFN in the serum of 177 SD patients and 28 HCs. Each individual was given a score relative to the mean HC expression of all modules in module M1.2. This correlated with Simoa IFN-α (Module M3.4 Spearman's rank r = 0.84, p < 0.0001 Module M5.12 r = 0.74, p < 0.0001).There is a weaker association between serum IFN-α and the IFN transcriptomic response in Chaussabel modules M3.4 and M5.12 than between serum IFN-α and Chaussabel module M1.2.

(F) IFN-α concentrations are not associated with symptom-based subgroups (Tarn et al. Lancet Rheumatology 2019), In the UKPSSR cohort four subgroups: Low symptom burden (LSB), high symptom burden (HSB), dryness dominant with fatigue (DDF), and pain dominant with fatigue (PDF). NA = not applicable Kruskal-Wallis p = 0.2942 (G, H) IFN-α concentrations are not associated with anxiety or depression in the UKPSSR.

(I) Transcriptomic expression of *SSB* and *TROVE2* was measured using a whole blood microarray in 177 SjD patients. There was no correlation seen with serum IFN-α (Spearman's rank r=0.03, p=0.7, r=0.01, p=0.9). Transcriptomic expression of *TRIM21* and association with serum IFN-α reproduced from figure 3B and shown alongside for comparison, (Spearman's rank r=0.53, p<0.0001).

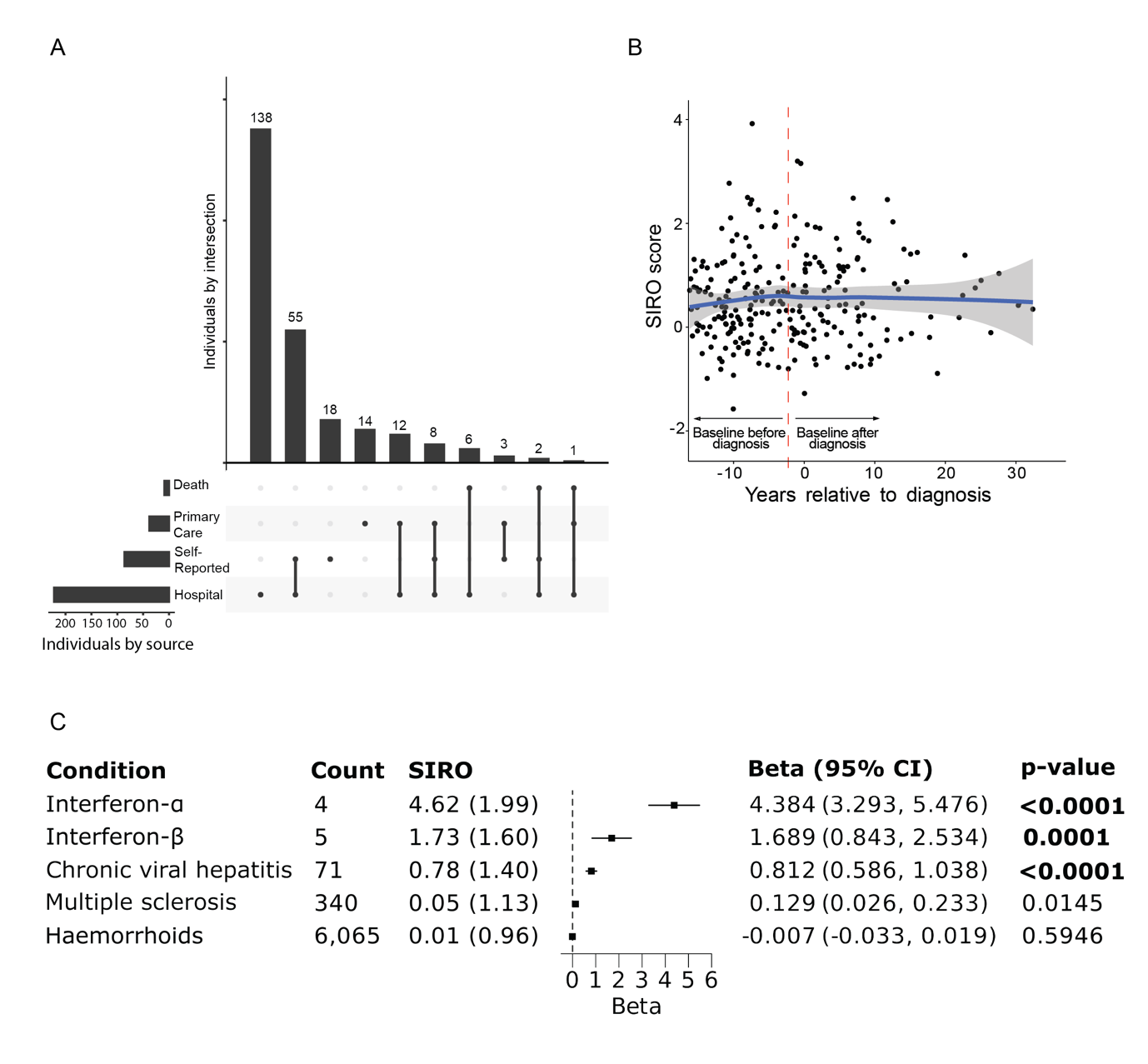

**Supplemental Figure 2. Sjögren Disease in UK Biobank**

(A) An Upset plot to demonstrate the sources of diagnosis for SjD cases in UK Biobank. There were 257 participants with SjD, which were mostly defined from hospital data alone (53.7%) or combined with other sources (32.7%), whereas 18 cases (7.0%) were self-reported only.

(B) SIRO scores did not significantly increase prior to the diagnosis of SjD in the UKB in linear regressions of SIRO by time before diagnosis (β per year = 0.02; 95% CI: -0.02, 0.06; p = 0.344), showing individual data points. Smooth curves with standard errors using the locally estimated scatter plot smoothing (LOESS) method.

(C) SIRO score in individuals in UK Biobank treated with recombinant type I interferon therapies, and SIRO scores of relevant diseases.

SIRO – Sjögren Interferon response in Olink.

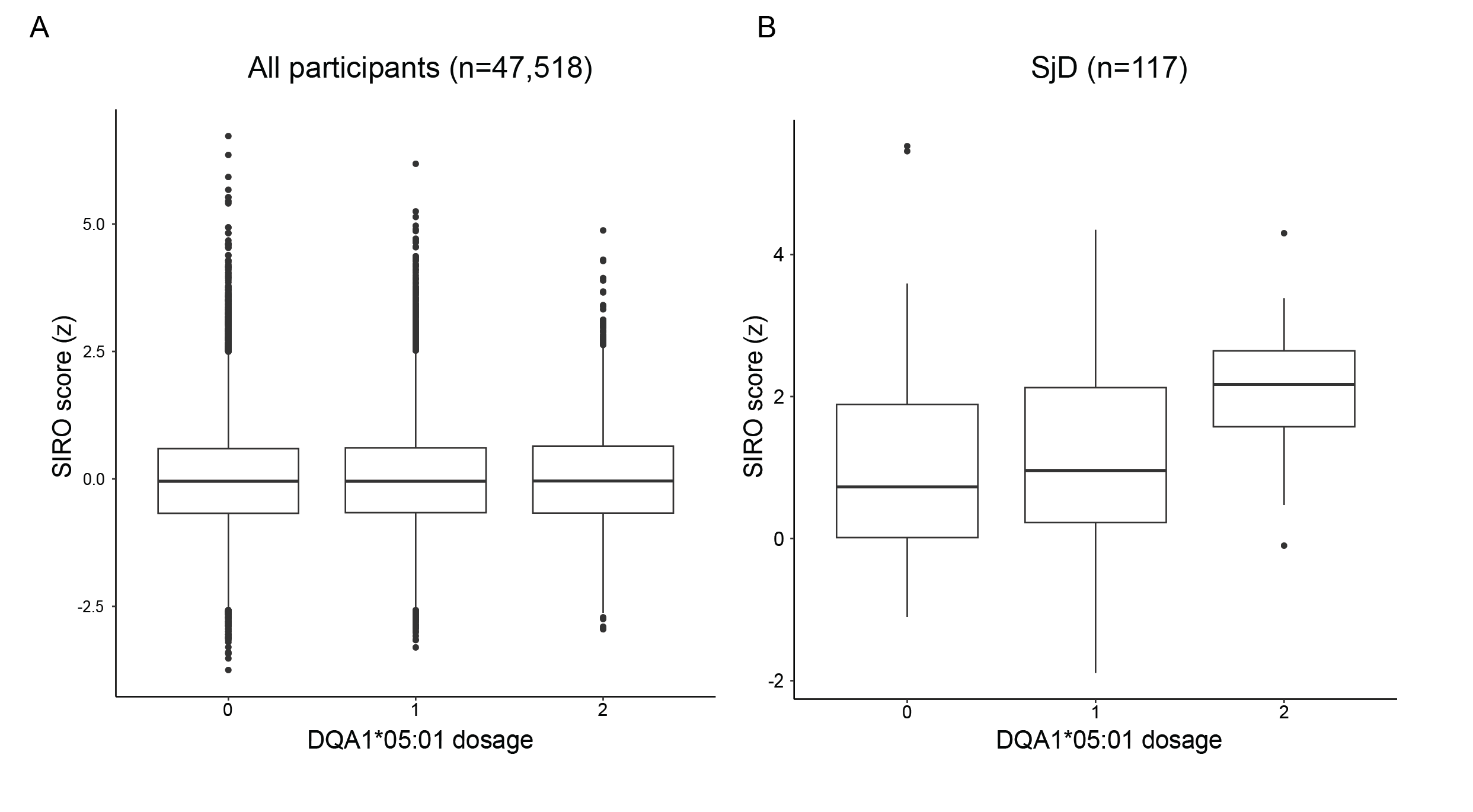

**Supplemental Figure 3. Oligoprotein interferon signature and HLA-DQA1*05:01 in SjD in UK Biobank.** The risk of SjD was significantly increased in carriers of ≥1 HLA-DQA1*05:01 allele as compared to non-carriers (OR=1.60; 95% CI: 1.45, 1.77; p<0.0001). (A) Carrying ≥1 HLA-DQA1*05:01 allele did not confer significantly higher SIRO scores in 47,518 participants with available data (beta=0.012; 95% CI: -0.006, 0.030; p=0.189) or 117 participants diagnosed with SjD before baseline (beta=0.075; 95% CI: -0.520, 0.671; p=0.802). (B) In people with SjD, however, carrying 2 HLA-DQA1*05:01 alleles significantly increased SIRO scores as compared to those with 0-1 alleles (beta=1.369; 95% CI: 0.188, 2.549; p=0.024).

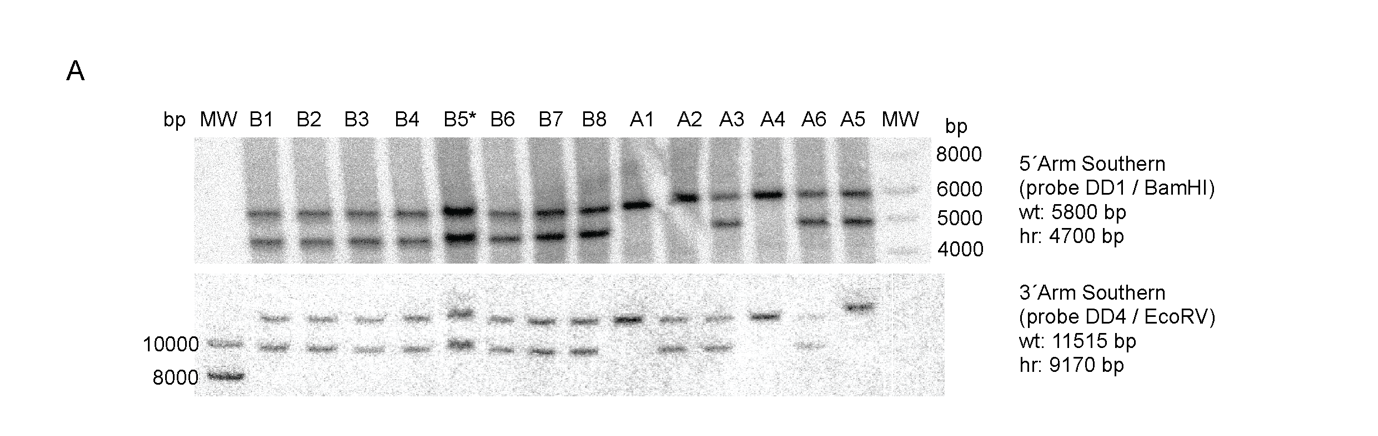

**Supplemental Figure 4. Generation of Rosa26-mIFNa4 transgenic mice**

(A) Rosa26-mIfn𝛼4 mice were generated by electroporation of the linearized targeting construct into Agouti JM8A1.N3 mouse embryonic stem cells resulting in 14 neomycin-resistant ES cell clones. Southern blot for identification of Agouti JM8A1.N3 mESC clones with genomic integration events involving the 5´ (upper) and 3´ (lower) arms of homology of the targeting construct is shown. Results were confirmed by long range PCR (not shown). Clone B5 was injected into C57BL/6N morulae and germline transmission was achieved to establish the Rosa26-mIFN𝛼4(“R26-mIFN𝛼4”) mouse line.

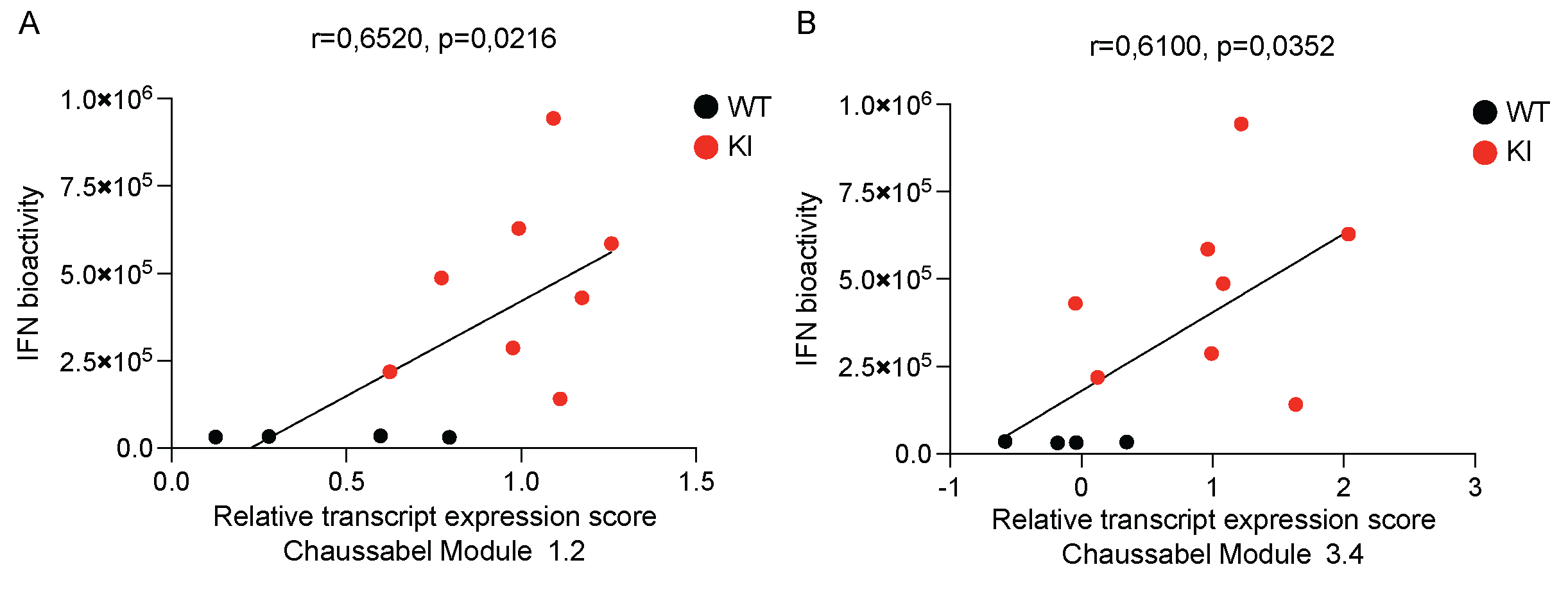

**Supplemental Figure 5. Serum IFN-α is associated with the type I IFN transcriptomic response in *Clec9a^KI/WT^R26Ifnα4^KI/WT^* mice.**

Transcript expression for the genes in the Chaussabel modules 1.2 and 3.4 was quantified. There was a significant relationship between Ifna and the relative expression of these interferon stimulated genes, a comparable result to the human analysis in Figure 1C.

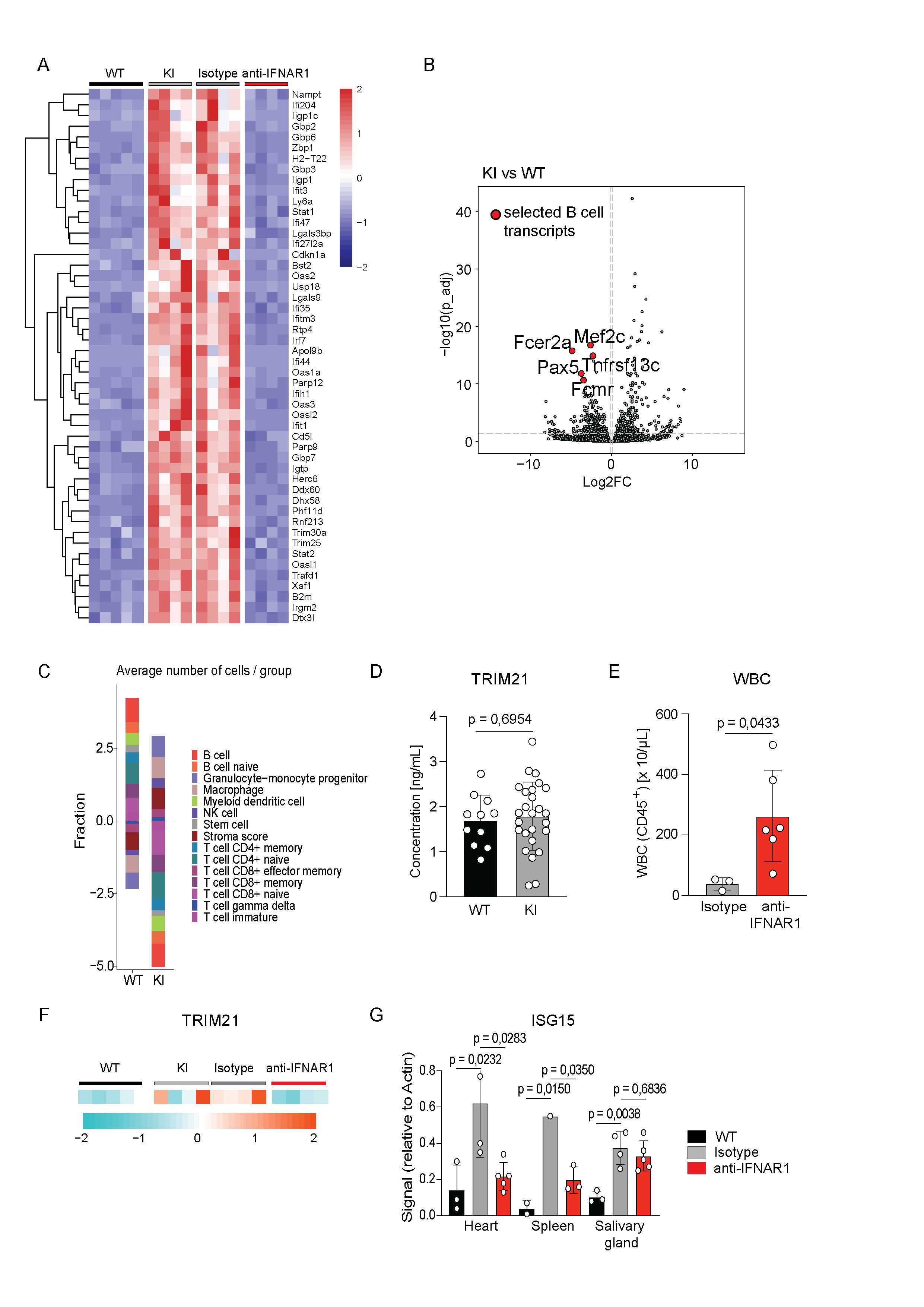

**Supplemental Figure 6. The role of chronic IFN-α4 elevation and IFNAR blockade in the induction of Sjögren disease specific immune alterations.**

A) Heatmap showing the top 50 genes differentially between WT (n=5), KI (n=4), Isotype (n=4) and anti-IFNAR1 treated (n=4) KI mice identified by two independent 3´RNA seq experiments on total RNA from splenocytes. Experiment 1 compared KI vs WT mice and experiment 2 Isotype-treated vs anti-IFNAR1-treated mice. All data are pooled for this analysis.

B) Volcano plot of all analysed genes in RNAseq experiment 1 comparing KI vs WT mice. B cell-specific transcripts (“red” dots) are significantly downregulated by chronic IFN-α4 overexpression (-log10 (p-adjusted) threshold = 1.3, Benjamini-Hochberg corrected p-value < 0.05).

C) Relative enrichment of cell type specific transcripts in whole-spleen RNAseq samples for estimation of immune cell type frequencies in WT and KI spleens.

D) Concentration of TRIM21 protein in sera of WT (n=11) and KI (n=26) mice quantified by ELISA.

E) WBC count (defined as DAPI^-^CD45^+^) in the whole blood of Isotype (n=3) and anti-IFNAR1 (n=6) treated KI mice quantified by flow cytometry.

F) Heatmap showing relative abundance of TRIM21 transcripts in splenocytes of WT (n=5), KI (n=4), Isotype (n=4) and anti-IFNAR1 treated (n=4) KI mice. Data extracted from the two RNAseq experiments shown in Figure S4A.

G) Quantification of ISG15 protein levels in heart, spleen and salivary glands of WT (n**>**2), KI (n**>**9), Isotype (n**>**1) and anti-IFNAR1 (n**>**3) treated KI mice.

Data are presented as a mean ± SD. Statistical significance was determined by Unpaired t-test or Mann-Whitney (D,E) or one-way ANOVA and Kruskal-Wallis test (G) depending on the data distribution.

A

B

C

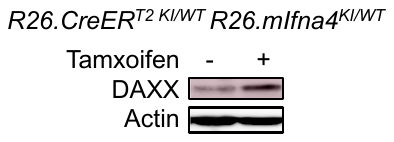

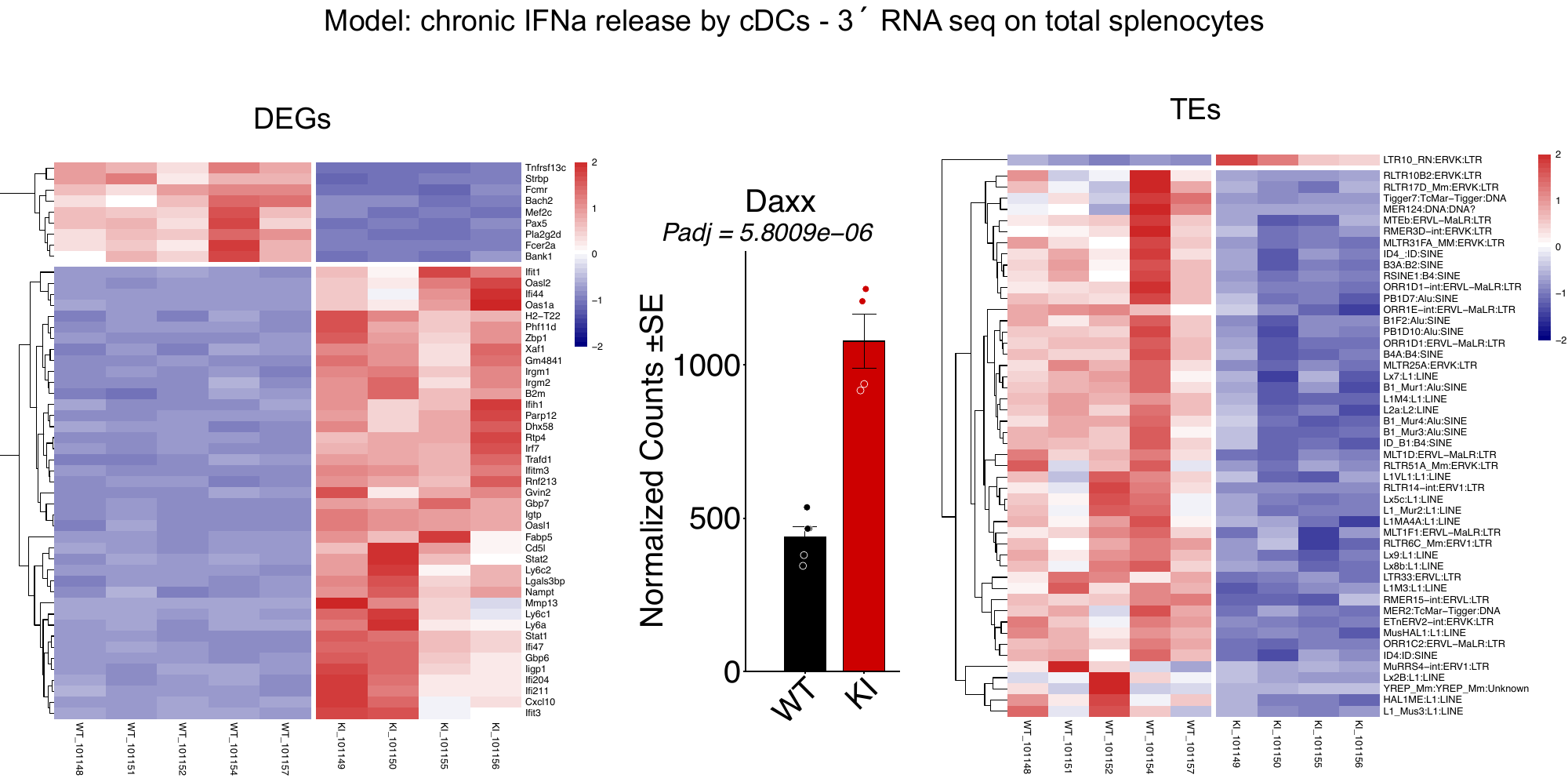

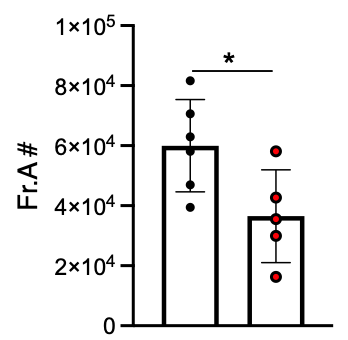

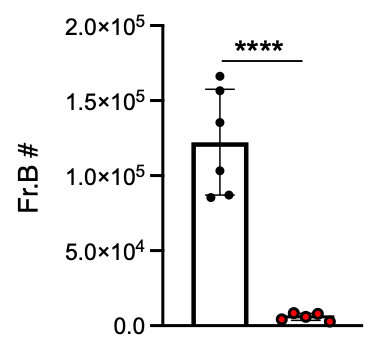

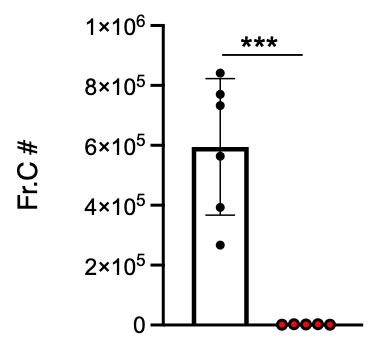

**Supplementary Figure 7. Chronic IFN-α exposure leads to upregulation of Daxx, associated with a block in B cell development at the pro-B cell stage**

(A) Transcript levels of *Daxx* in spleens of the indicated mice (RNAseq data).
(B) Daxx protein is induced 2 days after tamoxifen induced Cre-mediated activation of mIFNa4 in cultured splenic B cells.
(C) B cell developmental stages after Hardy et al., (PMID: 1827140) identified an almost complete block in B cell development from Fraction B containing predominantly late pro B cells. (black dots = WT, red dots = KI)

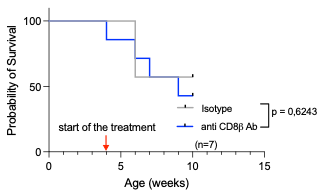

**Supplemental Figure 8. Depletion of CD8β -positive T cells in KI mice does not ameliorate autoimmunity.** *Clec9a.Cre^KI/WT^R26-mIfna4^KI/WT^* mice were treated once a week i.p. with 250 μg rat anti-mouse– CD8β antibody (Ab) (clone YTS156.7.7, anti- CD8β) or isotype-matched irrelevant rat-anti-Phyt1 (clone AFRC-MAC51, Isotype).

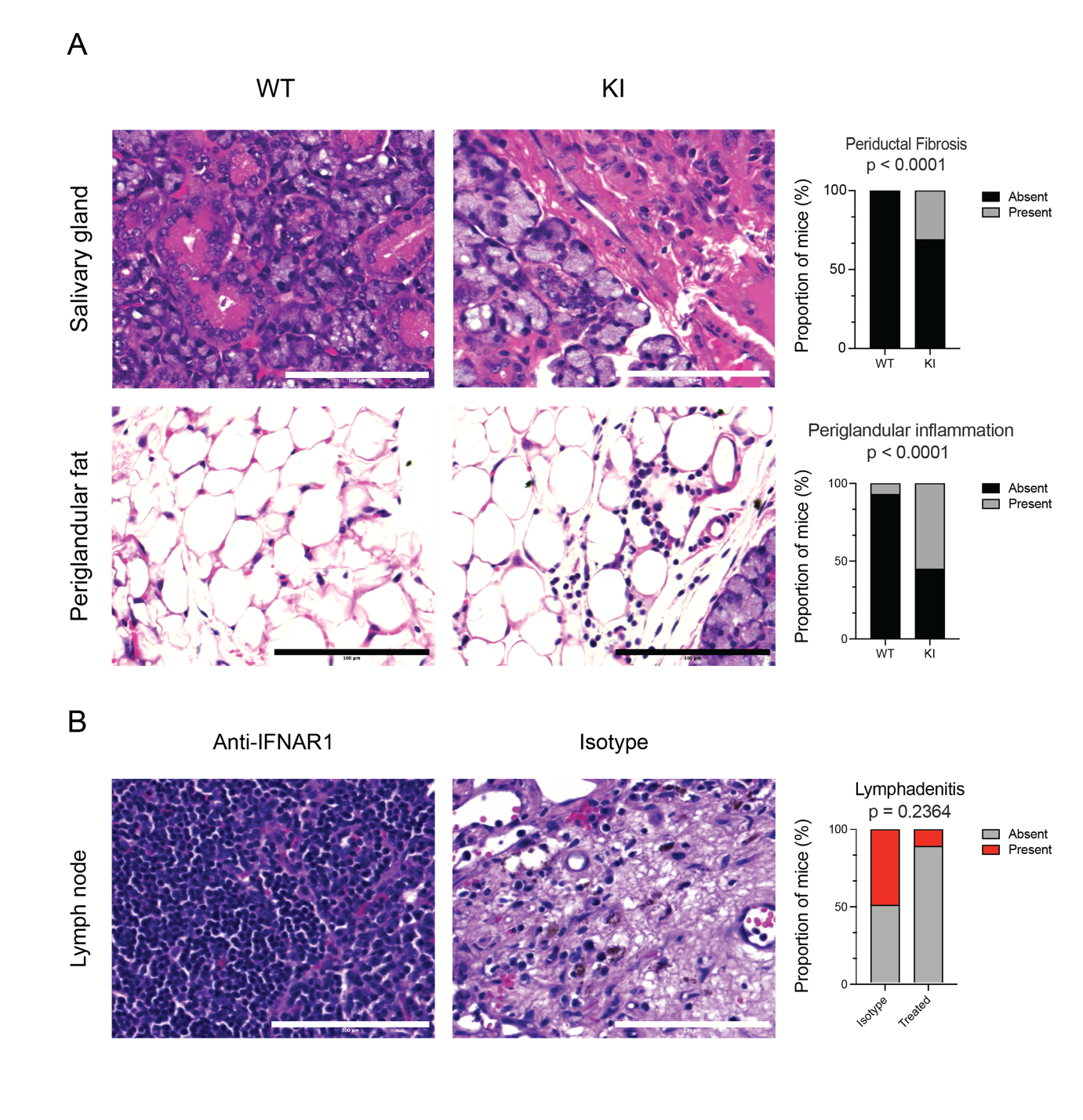

**Supplemental Figure 9. Chronic IFN-α4 elevation results in inflammation of salivary glands and periglandular fat**

(A) The Clec9a-R26Ifna4 mice have higher proportions of periductal fibrosis (WT 0% 0/14 vs. KI 11/27 29% p=0.0246) and lymphocytic infiltration of periglandular fat (WT 1/14 7.14% vs. 21/38 55.26% p=0.0017) upon expert pathological examination. (Two sided Fisher’s exact test performed in Prism Graphpad v9.0)

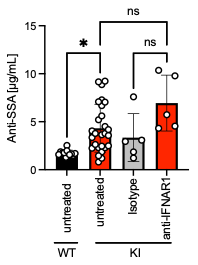

**Supplemental Figure 10.** **Circulating TRIM21 antibodies are not reduced by treatment with anti-IFNAR monoclonal antibody**. Concentration of TRIM21 antibody titres in isotype (n=5) and treated (n=5) mice, with untreated group (WT and KI), included for comparison.

(Note, baseline untreated groups include data points from figure 4J).

**Supplementary Table 1. Demographics of the UKPSSR cohort, Simoa substudy and Olink substudy**

| **Demographics** | **Whole cohort (n=1097)** | **Simoa cohort (n=177)** | **AAB Panel Cohort (n = 140)** | **OLINK Cohort**  **(n = 39)** |
| --- | --- | --- | --- | --- |
| **Age (median, (IQR))** | 59 (49, 67) | 57.5, (46,65) | 58 (46, 64.5) | 61 (58.5, 63.5) |
| **Sex Female %** | 992/1097 (90%) | 163/177 (92%) | 130/140 (93%) | 37/39 |
| **Ever a smoker %**  **(*n missing = 431, 40%)** | 210/1097(19%) | 39/177 (22%) | 28/140 (20%) | 12/39 |
| **BMI (median, (IQR))** | 25.7 (22.8, 29.5 | 25.9, (23.0, 29.1) | 26.0 (23.0, 28.9) | 27.3 (23.7, 29.4) |
| **SjD duration in years (median, (IQR))** | 4, (1, 8) | 4, (2, 8) | 5 (2, 8) | 4 (1, 7) |
| **Race white %** | 994/1097 (91%) | 162/177 (91%) | 125/140 (90%) | 37/39 (95%) |
| **Race non white %** | 70/1097(9%) | 15/177 (9%) | 15/140 (11%) | 2/39 (5%) |
| **ANA antibodies % (*78% Unknown)** | Unk: 856, Pos: 153, Neg: 88 | Unk: 146, Pos: 17, Neg: 14 | Unk: 110, Pos: 16, Neg: 14 | Unk: 31, Neg: 7, Pos: 1/39 |
| **Ro Antibodies %** | 855/1097  (88%) | 155/177  (88%) | 121/140 (86%) | 35/39 (90%) |
| **La Antibodies %** | 599/1097  (55%) | 106/177  6 Unknown  (60%) | 84/140 (60%)  Unk: 5 | 35/39  (90%) |
| **Immunosuppressive treatment at baseline %** | 751/1097  (68%) | 121/177  (68%) | 93/140 (66%) | 24/39  (62%) |
| **Baseline Disease activity ESSDAI (median, (IQR))** | 3, (1,7) | 3, (1,6) | 2 (0, 6) | 2 (0, 5.5) |
| **Baseline symptom score ESSPRI (median, (IQR))** | 5.7, (4, 7) | 5.5, (2.33, 7.33) | 5.33 (2.33, 7.33) | 2.7 (1.83, 7) |

**Supplementary Table 2. Elevated IFN-α in individuals with Sjögren disease is associated with a distinct immune endotype characterised by white blood cell cytopenias, hypergammaglobulinaemia and autoantibody production.**

|  | **IFN-α elevated >0.279pg/mL n=108 (61%)** | | **IFN-α normal**  **<0.279pg/mL n=69 (39%)** | |  |  |
| --- | --- | --- | --- | --- | --- | --- |
|  | **Median (IQR)** | **% Abnormal** | **Median (IQR)** | **% Abnormal** | **p Value**  **Fisher’s exact** | **Holm Sidak correction** |
| **Leukopenia** | 4.61  (1.0-5.6) | 43.0 | 5.6  (4.6-7.2) | 23.0 | 0.006 | 0.0355 |
| **Lymphopenia** | 1.31  (1.04-1.6) | 69.0 | 1.56  (1.2-1.96) | 50.0 | 0.0065 | 0.0355 |
| **IgG** | 17  (14-24) | 58.3 | 11.5  (9.6-16.6) | 27.5 | <0.0001 | 0.0011 |
| **IgA** | 3.2  (2.42-4.1) | 55.6 | 2.48  (1.6-3.5) | 37.7 | 0.002 | 0.0159 |
| **IgM** | 1.2  (0.76-1.62) | 6.00 | 1.19  (0.9-1.8) | 10.1 | 0.439 | 0.737 |
| **C3** | 1.16  (1.05-1.31) | 4.60 | 1.21  (1.01-1.37) | 0.00 | 0.359 | 0.737 |
| **C4** | 0.21  (0.17-0.27) | 15.7 | 0.22  (0.17-0.28) | 11.6 | 0.684 | 0.737 |
| ***Antibodies*** | **% seropositive** | | **% seropositive** | |  | |
| **ANA** | 50.0 | | 25.0 | | 0.0044 | 0.030 |
| **Ro/SSA** | 97.7 | | 55.3 | | <0.0001 | 0.0011 |
| **La** | 75.6 | | 40.4 | | <0.0001 | 0.0011 |
| **Ro52/TRIM21** | 68.3 | | 31.7 | | <0.007 | 0.0355 |

Units Leukopenia, Lymphopenia = cells *10^9^/L , IgG, IgA, IgM, C3, C4 = g/L.

**Supplementary Table 3. ESSDAI domains in the UKPSSR cohort**

|  | **Whole cohort** | **IFN-α normal <0.279pg/mL** | **IFN-α elevated >0.279pg/mL** | **p-value (with adjusted Holm-Sidak)** |
| --- | --- | --- | --- | --- |
| **ESSDAI Constitutional** | 20.2 | 18.8 | 24.1 | 0.53, 1 |
| **ESSDAI Lymphadenopathy/lymphoma** | 5.6 | 2.9 | 7.4 | 0.35, 1 |
| **ESSDAI Glandular** | 16.0 | 18.8 | 15.7 | 0.7409, 1 |
| **ESSDAI Articular** | 30.2 | 26.1 | 28.7 | 0.84, 1 |
| **ESSDAI Cutaneous** | 6.6 | 4.3 | 5.6 | 0.99, 1 |
| **ESSDAI Pulmonary** | 9.5 | 8.7 | 6.5 | 0.80, 1 |
| **ESSDAI Renal** | 2.4 | 2.9 | 3.7 | 1, 1 |
| **ESSDAI Muscular** | 2.0 | 1.4 | 0.9 | 1, 1 |
| **ESSDAI PNS** | 5.2 | 1.4 | 3.7 | 0.68, 1 |
| **ESSDAI CNS** | 0.7 | 1.4 | 0.0 | 0.82, 1 |
| **ESSDAI Haematological** | 13.9 | 10.1 | 14.8 | 0.50, 1 |
| **ESSDAI Biologic** | 40.4 | 30.4 | 45.4 | 0.0027, 0.06 |

**Supplementary Table 4. Demographics of the UK Biobank cohort and UKB-PPP substudy**

| **Characteristic** | **UKB-PPP**  **(n=** **53,014)** | **SIRO available** | | **SIRO not available** | |
| --- | --- | --- | --- | --- | --- |
|  |  | **SjD**  **(n=** **257)** | **No SjD**  **(n=** **47,606)** | **SjD**  **(n=25)** | **No SjD**  **(n=** **5,126)** |
| **Age at recruitment, mean (SD)** | 56.8 (8.2) | 58.6 (7.5) | 56.8 (8.2) | 60.2 (6.4) | 56.8 (8.2) |
| **Female sex, n (%)** | 28,580 (53.9) | 232 (90.3) | 25,610 (53.8) | 21 (84.0) | 2,717 (53.0) |
| **Ethnic background, n (%)** | | | | | |
| **White** | 49,435 (93.2) | 234 (91.1) | 44,455 (93.4) | 25 (100) | 4,721 (92.1) |
| **Black/Black British** | 1,217 (2.3) | 10 (3.9) | 1,074 (2.3) | 0 (0) | 133 (2.6) |
| **Asian/Asian British** | 986 (1.9) | 7 (2.7) | 857 (1.8) | 0 (0) | 122 (2.4) |
| **Other** | 619 (1.2) | 3 (1.2) | 554 (1.2) | 0 (0) | 62 (1.2) |
| **Mixed** | 349 (0.7) | 2 (0.8) | 308 (0.6) | 0 (0) | 39 (0.8) |
| **Chinese** | 148 (0.3) | 0 (0) | 132 (0.3) | 0 (0) | 16 (0.3) |
| **Missing** | 260 (0.5) | 1 (0.4) | 226 (0.5) | 0 (0) | 33 (0.6) |
| **Age at diagnosis, mean (SD)** | NA | 58.8 (12.5) | NA | 63.1 (10.2) | NA |

UKB-PPP subset excludes participants from the COVID-19 repeat imaging substudy. Abbreviations: SD, standard deviation; SjD, Sjogren disease.

Tarn, J.R., N. Howard-Tripp, D.W. Lendrem, X. Mariette, A. Saraux, V. Devauchelle-Pensec, R. Seror, A.J. Skelton, K. James, P. McMeekin, S. Al-Ali, K.L. Hackett, B.C. Lendrem, B. Hargreaves, J. Casement, S. Mitchell, S.J. Bowman, E. Price, C.T. Pease, P. Emery, P. Lanyon, J. Hunter, M. Gupta, M. Bombardieri, N. Sutcliffe, C. Pitzalis, J. McLaren, A. Cooper, M. Regan, I. Giles, D. Isenberg, V. Saravanan, D. Coady, B. Dasgupta, N. McHugh, S. Young-Min, R. Moots, N. Gendi, M. Akil, B. Griffiths, S.J.A. Johnsen, K.B. Norheim, R. Omdal, D. Stocken, C. Everett, C. Fernandez, J.D. Isaacs, J.-E. Gottenberg, W.-F. Ng, V. Devauchelle-Pensec, P. Dieude, J.J. Dubost, A.-L. Fauchais, V. Goeb, E. Hachulla, C. Larroche, V. Le Guern, J. Morel, A. Perdriger, X. Puéchal, S. Rist, D. Sen, J. Sibilia, O. Vittecoq, J. Benessiano, S. Tubiana, K. Inamo, S. Gaete, D. Batouche, D. Molinari, M. Randrianandrasana, I. Pane, A. Abbe, G. Baron, P. Ravaud, J.-E. Gottenberg, P. Ravaud, X. Puéchal, V. Le Guern, J. Sibilia, C. Larroche, A. Saraux, V. Devauchelle-Pensec, J. Morel, G. Hayem, P. Hatron, A. Perdriger, D. Sene, C. Zarnitsky, D. Batouche, V. Furlan, J. Benessiano, E. Perrodeau, R. Seror, X. Mariette, S. Brown, N.C. Navarro, C. Pitzalis, P. Emery, S. Pavitt, J. Gray, C. Hulme, F. Hall, R. Busch, P. Smith, L. Dawson, M. Bombardieri, W.F. Ng, C. Pease, E. Price, N. Sutcliffe, C. Woods, S. Ruddock, C. Everett, C. Reynolds, E. Skinner, A. Poveda-Gallego, J. Rout, I. Macleod, S. Rauz, S. Bowman, W.-F. Ng, S.J. Bowman, B. Griffiths, F. Hall, E.C. Bacaba, H. Frankland, R. Moots, K. Chadravarty, S. Lamabadusuriya, M. Bombardieri, C. Pitzalis, N. Sutcliffe, C. Breston, N. Gendi, K. Culfear, C. Riddell, J. Hamburger, A. Richards, S. Rauz, S. Brailsford, J. Dasgin, J. Logan, D. Mulherin, J. Andrews, P. Emery, A. McManus, C. Pease, D. Pickles, A. Booth, M. Regan, J.K. Kin, A. Holt, T. Dimitroulas, L. Kadiki, D. Kaur, G. Kitas, A. Khan, T. Cosier, Panthakalam, K. Mintrim, M. Lloyd, L. Moore, E. Gordon, C. Lawson, M. Gupta, J. Hunter, L. Stirton, G. Ortiz, E. Price, S. Pelger, C. Gorman, B. Hans, G. Clunie, S. Lane, G. Rose, S. Cuckow, M. Batley, R. Einosas, S. Knight, D. Symmons, B. Jones, A. Carr, S. Edgar, F. Figuereido, H. Foggo, D. Lendrem, I. Macleod, S. Mitchell, C. Downie, J. Tarn, J. Locke, S. Al-Ali, S. Legg, K. Mirza, B. Hargreaves, L. Hetherington, A. Jones, P. Lanyon, A. Muir, P. White, S. Young-Min, S. Pugmire, S. Vadivelu, A. Cooper, M. Watkins, A. Field, S. Kaye, D. Mewar, P. Medcalf, P. Tomlinson, D. Whiteside, N. McHugh, J. Pauling, J. James, A. Dowden, M. Akil, J. McDermott, O. Godia, D. Coady, E. Kidd, L. Palmer, C. Li, S. Bartrum, D. Mead, B. Dasgupta, V. Katsande, P. Long, E. Vermaak, J. Turner, U. Chandra, K. Mackay, S. Fedele, A. Ferenkeh-Koroma, I. Giles, D. Isenberg, H. Maconnell, N. Ahwiren, S. Porter, P. Allcoa, and J. McLaren. 2019. Symptom-based stratification of patients with primary Sjögren's syndrome: multi-dimensional characterisation of international observational cohorts and reanalyses of randomised clinical trials. *The Lancet Rheumatology* 1:e85-e94.

Trutschel, D., P. Bost, X. Mariette, V. Bondet, A. Llibre, C. Posseme, B. Charbit, C.W. Thorball, R. Jonsson, C.J. Lessard, R. Felten, W.F. Ng, L. Chatenoud, H. Dumortier, J. Sibilia, J. Fellay, K.A. Brokstad, S. Appel, J.R. Tarn, L. Quintana-Murci, M. Mingueneau, N. Meyer, D. Duffy, B. Schwikowski, J.E. Gottenberg, A.s.i. Milieu Interieur Consortium, and N. Consortium. 2022. Variability of Primary Sjogren's Syndrome Is Driven by Interferon-alpha and Interferon-alpha Blood Levels Are Associated With the Class II HLA-DQ Locus. *Arthritis Rheumatol* 74:1991-2002.

Uzé, G., S. Di Marco, E. Mouchel-Vielh, D. Monneron, M.T. Bandu, M.A. Horisberger, A. Dorques, G. Lutfalla, and K.E. Mogensen. 1994. Domains of interaction between alpha interferon and its receptor components. *J Mol Biol* 243:245-257.
